# The dynamic fitness landscape of ageing haematopoiesis through clonal competition

**DOI:** 10.1101/2024.04.16.589764

**Authors:** Nathaniel Mon Père, Francesco Terenzi, Benjamin Werner

## Abstract

Clonal haematopoiesis (CH) – the existence of large mutant clones in blood – is a prime example of somatic evolution. Yet how evolution shapes CH with age remains to be understood. Here, we show that clonal competition can explain the complex dynamics observed in vivo. In this paradigm, numerous fit clones continually appear and compete, driving an evolving fitness landscape of stem cells. This naturally explains shrinking expanded clones, varying driver efficacy across individuals, and transitions of site frequency spectra with age. Inferences of evolutionary parameters from variant trajectories and site frequency spectra converge to nearly identical estimates of a non-exponential fitness distribution with mean 0.08, and an arrival rate of 2-20 advantageous clones per year. Inferring innate fitnesses from single trajectories, we find that 80% of the variance on identical mutations is explained by clonal competition, with fitness estimates of most common drivers between 0.14 and 0.18. Strikingly, we find clones with much higher fitness to occur only later in life, with an arrival time distribution well-described by a multi-step model of clonal evolution. Overall, a quantitative clonal competition model predicts many aspects of ageing haematopoiesis and allows a personalized identification of high-risk clones potentially important for patient stratification.

## Introduction

Genetic heterogeneity pervades somatic tissues at all stages of human life [1–5]. It emerges from ubiquitous mutational processes and subsequent expansion of novel clones by selection and random drift [6–10]. This phenomenon has been well-characterised in blood, where largely expanded clones are common late in life, a condition termed clonal haematopoiesis (CH) [11–16]. Still, obtaining a detailed quantitative description of the evolutionary forces of CH and how these change with age, mutational load, and tissue regulation remains challenging.

The simplest evolutionary models of CH assume that positively selected somatic mutations in haematopoietic stem cells (HSCs) are rare and their fitnesses are context-independent [7, 8, 17]. However, recent observations suggest a more complex scenario [5, 15, 16, 18]. Clone frequencies measured at multiple time points exhibit saturating growth curves despite possessing positive fitness [15]. While by itself such behaviour is consistent with a size-constrained HSC population [19, 20], other observations remain unexplained: clones typically saturate below fixation, many clone trajectories decrease late in life, and the average HSC fitness appears to decrease with age [15, 16]. Stratifying inter-individually further complicates this picture, as observable fitnesses of identical genetic variants vary greatly across donors [7, 15, 17, 21].

Here we show that these phenomena naturally emerge in an evolutionary model of clonal competition in which multiple clones of differing fitness consistently appear, compete against a unique collection of clones, and change the fitness landscape over time, in a way that is highly stochastic between individuals (Figure 1A). The clonal competition model generates testable longitudinal predictions of clone trajectories and variant frequency distributions that are measurable from *in vivo* observations.

**Figure 1.**
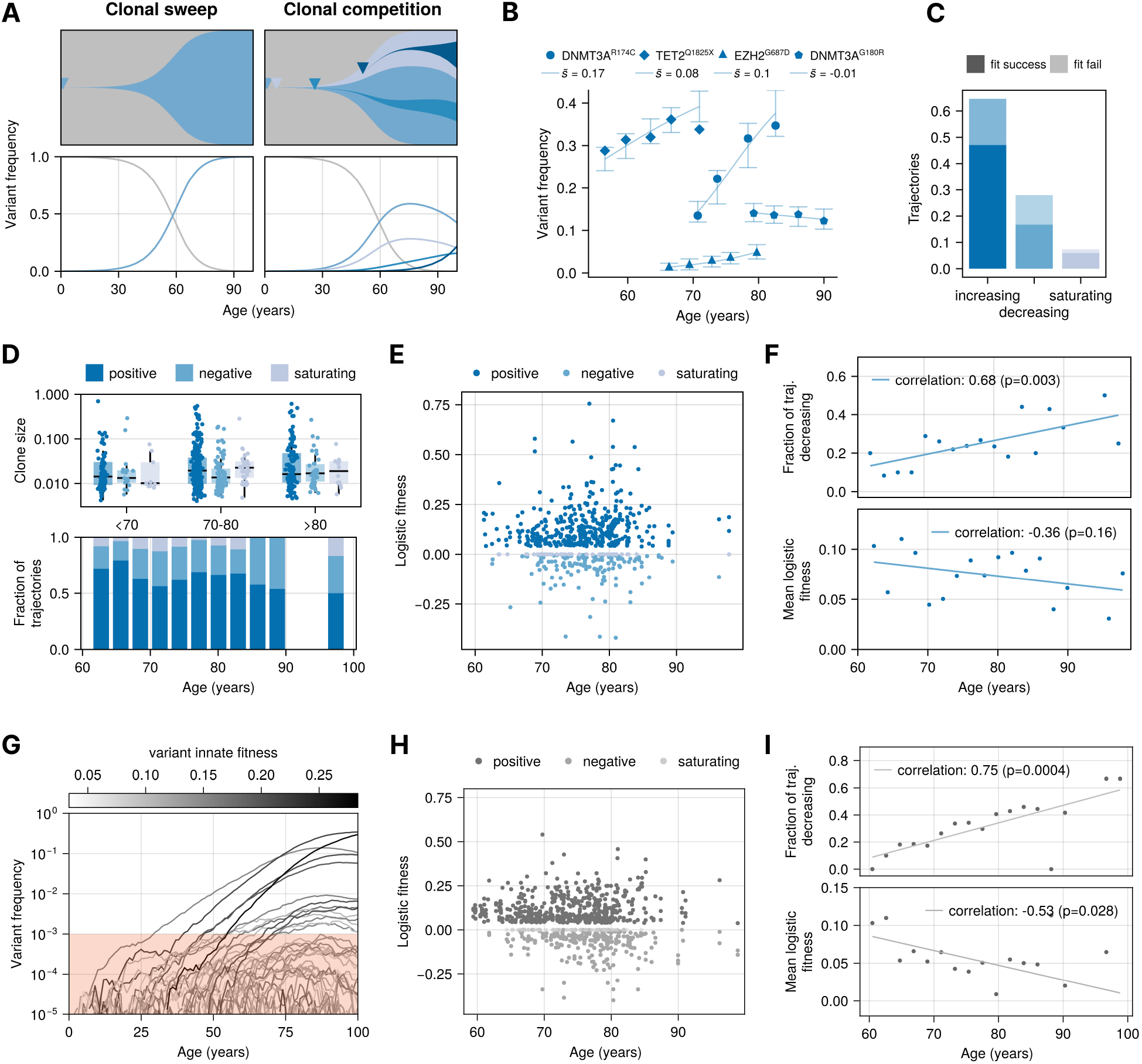
Changing fitness landscape of human haematopoiesis with age. **A)** Schematic of the expected trajectories of CH as clonal sweeps (fixation) or patterns of clonal interference. **B)** Variant allele frequency trajectories of CH variants with age from donor data (blue symbols), logistic fits (blue lines), and 95% confidence intervals obtained from maximum likelihood estimation. **C)** Fraction of increasing, decreasing, and saturating logistic fitnesses over 697 fitted variant trajectories. A successful fit is defined as all data points falling within the MLE-estimated 95% confidence interval. **D)** Clone sizes and trajectory composition with age. **E)** Logistic fitness coefficients *s*_log_ obtained from maximum likelihood estimation on all 697 variant trajectories. **F)** The fraction of decreasing trajectories increases with age (Spearman correlation = 0.68, *p* = 0.003). Mean logistic fitness decreases with age (Spearman correlation = -0.36, *p* = 0.16). **G)** Realisation of a stochastic simulation of CH from Equation (1). **H)** Logistic fitness coefficients *s*_log_ measured from simulations of CH with parameters *η* = 0.11, *σ* = 0.032, and *µ* = 5. **I)** The fraction of decreasing trajectories also increases with age in stochastic simulations (Spearman correlation = 0.75, *p* = 0.0004) and mean fitness decreases with age (Spearman correlation = -0.53, *p* = 0.028).

## Results

### Trajectories of clonal haematopoiesis suggest a dynamic fitness landscape in HSCs

To quantify the changing haematopoietic fitness landscape with age, we first revisited longitudinal measurements obtained by Fabre et al. [15] of variant allele frequencies of 56 known driver mutations across 385 individuals. This resulted in 697 variant trajectories with 13 years median follow-up time (Figure 1B). We evaluated the selection strength for each variant by fitting its change in frequency over time to a logistic differential growth function *dx*(*t*)/*dt* = *s*_log_*x*(*t*)(1 − *x*(*t*)), where *x*(*t*) denotes the frequency of a variant at age *t*. The logistic fitness coefficient *s*_log_ is either positive for increasing, negative for decreasing, or zero for saturating trajectories. We obtained best-fit fitness coefficients by implementing a maximum likelihood estimator that assumes the observed variant frequencies to be binomially sampled from unobserved true frequencies (Figure 1B, SI 1). This resulted in a distribution of logistic fitnesses for individuals ranging from 62 to 98 years of age (Figure 1E). Overall, 67% of all successfully fitted trajectories were increasing, 24% decreasing, and 8% saturating (Figure 1C-D). Similar frequencies were recently reported by Van Zeventer et al. in a set of 1,642 mutations across 1,573 individuals [21]. After stratifying trajectories by age, the relative proportion of decreasing trajectories increased from 18% at age 63 to 41% at age 88 (Spearman correlation=0.68, p=0.003), and the mean logistic fitness decreased with age (Spearman correlation=-0.36, p=0.16), (Figure 1F). Since variants in a population of approximately 10^5^ HSCs are unlikely to have reached detectable frequencies of 0.01 or higher solely by random drift [7, 8], those with decreasing trajectories must have been under positive selection in the past. This suggests an emerging dynamic HSC fitness landscape in which observable clone fitness can change with age.

### Clonal competition predicts varying CH trajectories within and across individuals

The logistic growth model allows quantification of a clone’s observable selection strength in a brief time window through its fitness parameter *s*_log_. However, mathematically this model describes the expected growth of only a single clone in a background population of static fitness [20]. We extended the single-clone model to account for the effects of a shared carrying capacity on trajectories of multiple coexisting clones in a context-dependent fitness landscape. Here, new clones arise at random times according to a Poisson process with constant rate *µ*. Each clone is assigned an innate fitness randomly drawn from a Gamma distribution, a flexible choice that contains many families of probability distributions as limiting cases. We show that the fully stochastic birth-death process of an arbitrary number of clones obeys a set of stochastic differential equations given by

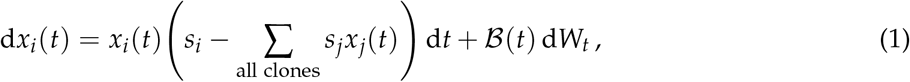

where *x*_*i*_(*t*) denotes the frequency of clone *i* at time *t*, and ℬ(*t*) is a fluctuation term with Gaussian noise d*W*_*t*_ (see SI 2.1). We denote *s*_*i*_ the innate fitness of clone *i*, defined as the increase of the clone’s proliferation rate relative to the wild type in the absence of competition: *s*_*i*_ = rate_*i*_ − rate_wt_. The deterministic part of Equation (1) describes a competitive Lotka-Volterra equation [22], however here the number of equations increases as clones emerge. We confirmed its validity with agent-based simulations of the underlying birth-death process using the Gillespie algorithm [23] (SI 4.2, Figure SI 1E). Equation (1) introduces an important distinction between a clone’s innate fitness *s*_*i*_ and its observable growth rate, measurable via sequencing through its change in clonal frequency over time. In fact, because the latter depends on the presence of all other clones (*s*_*i*_ − ∑_*j*_ *x*_*j*_*s*_*j*_), a clone with positive innate fitness can become negatively selected if this fitness is surpassed by the increasing average fitness in the population – that is if *s*_*i*_ < ∑_*j*_ *x*_*j*_*s*_*j*_ (Figure 1A, G). This becomes more likely with age as more fit clones inhabit the system. Consequently, as the clonal composition of the stem cell population grows, the effective fitnesses (i.e. observable growth rates) of individual clones become increasingly dynamic.

Using Equation (1) we simulated clone trajectories in virtual individuals that matched the age and frequency of clones in the data. We then applied the same maximum likelihood estimation to obtain the predicted age-stratified distribution of fitted logistic fitness coefficients *s*_log_ from simulated trajectories (Figure 1H). In accordance with the data, simulations show the emergence of decreasing variant trajectories with an increasing abundance with age (Spearman correlation = 0.75, *p* = 0.0004), while the average logistic fitness across all variants decreases with age (Spearman correlation = - 0.53, *p* = 0.028, Figure 1I). The model furthermore predicts that the measurable logistic fitness of a variant varies strongly with its time of occurrence, the individual’s age at measurement, and the unique landscape of co-occurring mutations (Figure SI 1F). This provides a natural explanation for the observed variation of fitnesses of identical CH driver mutations across individuals [15, 17, 21]. Together, these findings suggest that clonal competition could underlie the dynamic and stochastic nature of observed clone trajectories.

### Characteristic transitions of site frequency spectra with age reflect HSC population dynamics

To further probe the consequences of clonal competition, we quantified HSC genetic heterogeneity using the site frequency spectrum (SFS) – the distribution of prevalences of genetic variants within a population. The SFS is typically dominated by neutral mutations which arise far more frequently than positively selected variants. However, neutral mutations hitchhike on expanding fit clones [24], suggesting that a varying fitness landscape within the HSC population may change the SFS with age (Figure 2A).

**Figure 2.**
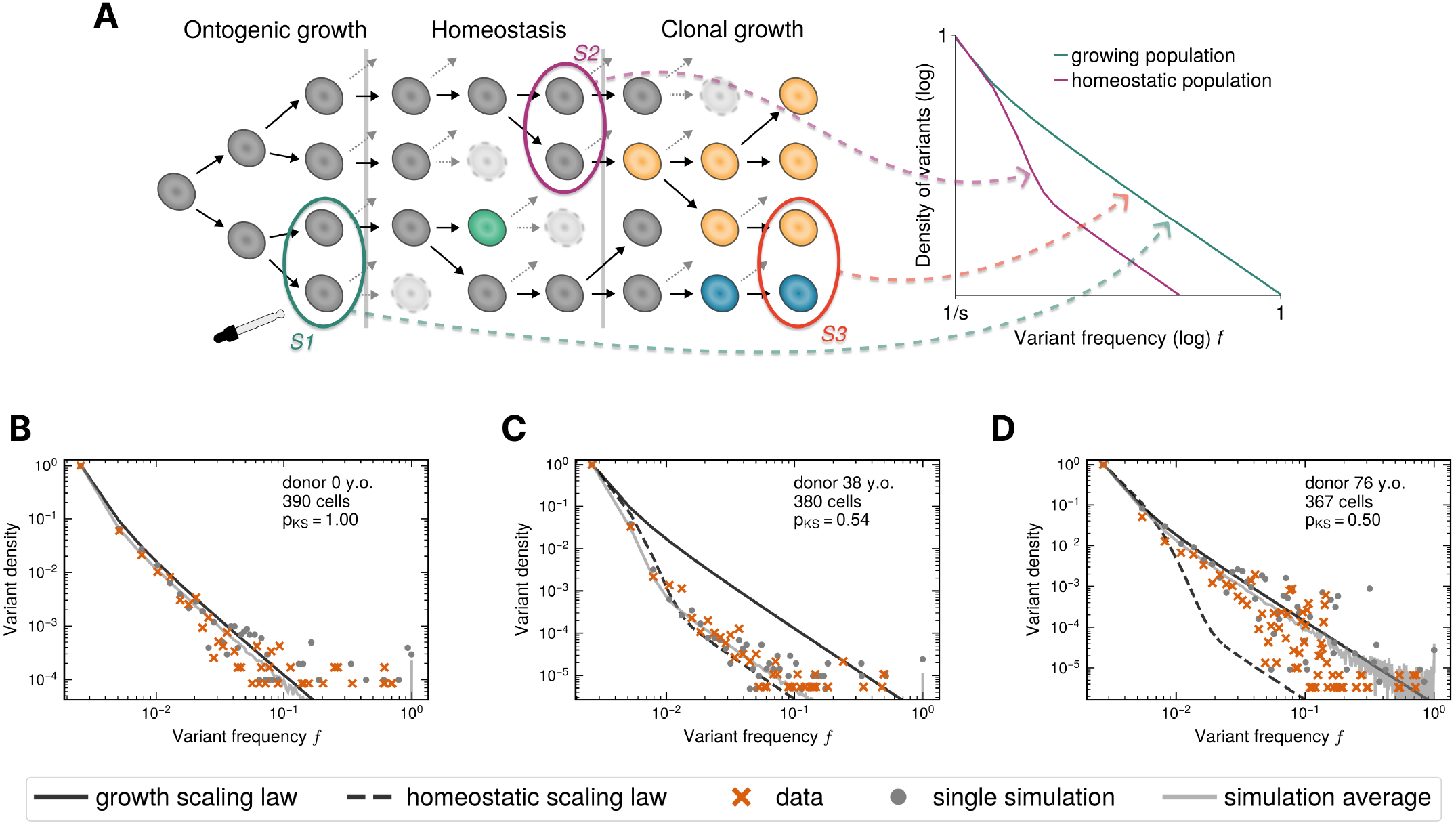
Transitions of HSC site frequency spectra with age. **A)** Schematic of the expected scaling shift of the SFS under ontogenic growth, homeostatisis, and clonal expansion of HSCs. Theory predicts three distinct phases of the SFS with age in the presence of clonal competition. **B)** The SFS of a newborn reconstructed from 390 whole genome sequenced single cells (orange crosses), a single realisation of the agent-based simulation (grey dots), averages over 500 simulations (grey line) and the expected scaling of an exponentially growing population (black line). **C)** SFS in a 38 year old donor constructed from 380 single HSCs. The SFS has shifted towards the expected scaling of a homeostatic population of constant size (black dashed line). **D)** SFS in a 76 year old donor constructed from 367 single HSCs. The SFS reverts to the scaling of a growing population at frequencies below < 0.1 (black line) and subclonal peaks appear.

To derive predictions for the expected SFS in the presence of clonal competition, we extended our model to include the accumulation of neutral variants in a population of 10^5^ HSCs through agent-based stochastic simulations (SI 4). From these simulations we observe two frequency domains of the SFS (Figure 2B-D, Figure SI 6). Variants at high frequencies (≳ 10^−1^) are sparse and highly stochastic across realisations. At young ages they are predominantly developmental in origin, whereas in the elderly they are a combination of developmental and positively-selected variants. In contrast, at lower frequencies (≲ 10^−1^) variant distributions follow power laws that emerge from the underlying HSC dynamics (Figure 2A). At birth the number of mutations at frequency *f* follows a 1/ *f* ^2^ power law. This is characteristic for expanding populations [25], and occurs here as a consequence of developmental growth. By early adulthood, the SFS resembles a wave asymptotically approaching a 1/ *f* power law, which is associated with a neutral homeostatic population of fixed size [10]. In the elderly our simulations suggest the SFS shifts again, with variants at low frequency reverting to a 1/ *f* ^2^ scaling due to the expansion of positively selected clones. Thus our stochastic simulations predict two transitions between three characteristic stages of the SFS in HSCs. The first reflects a deterministic change of the population level dynamics from a growing into a constant homeostatic HSC population. The second is driven by subclonal expansions and would be absent in a neutrally evolving population.

We tested these predictions in SFS reconstructed from single-cell whole genome sequencing data of HSCs published by Mitchell and colleagues [16] (Figure 2B-D). In brief, whole genome sequencing of 200 to 500 single HSCs in 9 healthy donors aged 0 to 81 allowed us to compute SFS with high resolution across the observed frequency spectrum. After adjusting for sampling, measured spectra of all ages agree well with stochastic simulations (Kolmogorov-Smirnov test *p*_*KS*_ > 5% for all SFS, Figure 2B-D, SI 6). Also noticeable is the increasing roughness at higher frequencies in the SFS in the elderly, suggesting unique evolutionary histories. Single realisations of our stochastic simulations reproduce the magnitude of these fluctuations that are absent in a population average (Figure SI 6). In summary, a clonal competition model precisely recovers the non-trivial dynamics of the SFS from newborns to octogenarians, capturing the changes in the genetic heterogeneity observed in clonal haematopoiesis at single-cell resolution.

### A Bayesian computation framework to estimate parameters of CH evolution

We next investigated the identifiability of the underlying evolutionary parameters of CH from *in vivo* data. The clonal competition model defines three parameters that determine the HSC fitness landscape: the mean *η* and standard deviation *σ* of the innate fitness distribution, and the arrival rate of advantageous clones *µ*. We additionally require the size of the adult HSC population *N* and the wild-type HSC inter-division time *τ*, which describe healthy haematopoiesis in the absence of selection.

To test what combinations of these parameters 𝒫_*i*_ = { *µ*_*i*_, *η*_*i*_, *σ*_*i*_, *N*_*i*_, *τ*_*i*_ } best describe the data, we implemented approximate Bayesian computation (ABC) inference schemes [26]. In brief, we randomly drew 65,000 parameter combinations 𝒫_*i*_ from uninformed uniform prior distributions. For each 𝒫_*i*_ we generated *in silico* predictions from our modelling and quantitatively compared them to donor data to obtain a metric distance. We then retained the 1% quantile of ranked distances to derive posterior distributions containing those 𝒫_*i*_ with predictions closest to the data (Figure 3A).

**Figure 3.**
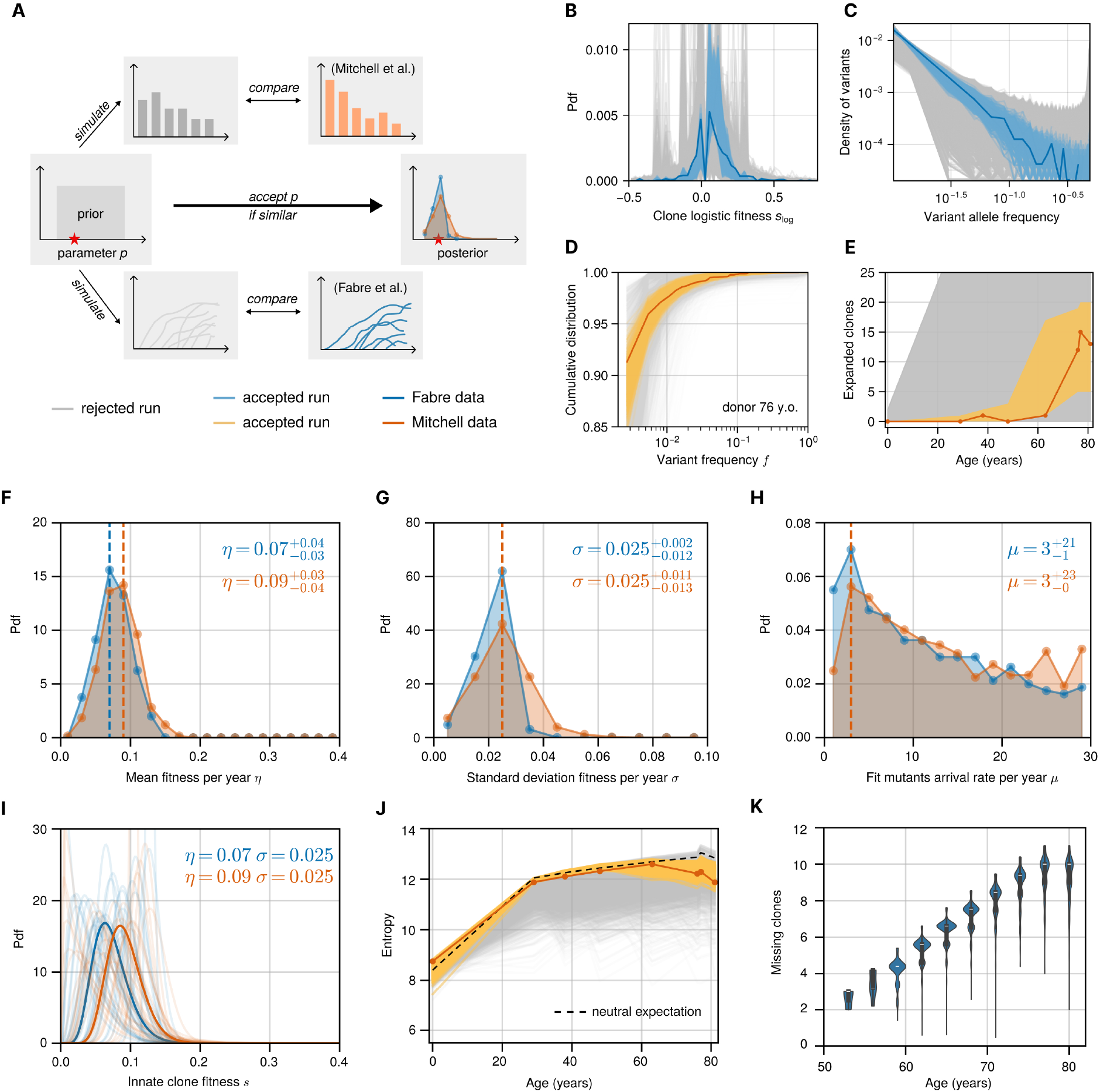
Inferences of the fitness distribution and arrival rate of advantageous clones in CH. **A)** Schematic of the approximate Bayesian computation (ABC) framework. **B)** Logistic fitness distribution *s*_log_ *in vivo* (dark blue lines), in ABC accepted runs (light blue lines), and in ABC rejected runs (grey lines). **C)** Clone size distribution *in vivo* (dark blue lines), in ABC accepted runs (light blue lines), and in ABC rejected runs (grey lines) runs. **D)** Cumulative SFS of the 76 year old donor (dark orange lines), in ABC accepted runs (light orange lines), and in ABC rejected runs (grey area). **E)** Number of visible clones at 1% resolution *in vivo* (orange dots), in ABC accepted runs (orange shaded area), and in ABC rejected runs (grey shaded area). **F-H)** Posterior ABC parameter estimates based on trajectories (blue) and SFS (orange) for the mean *η* and the standard deviation *σ* of the innate fitness distribution of advantageous variants, and the HSC population arrival rate *µ* per year of advantageous variants. **I)** Best estimates of the innate fitness distribution of advantageous variants based on trajectories (blue) and SFS (orange). Thin lines represent individual accepted runs. Thick lines correspond to the estimated mean and standard deviation. **J)** Entropy computed from the *in vivo* SFS (dark orange lines), simulated SFS of the ABC accepted runs (light orange lines)^1, 2^and simulated SFS of the ABC rejected runs (grey area). The neutral expectation (dashed black line) is the average over 100 SFS simulated with neutral dynamics. **K)** The expanded clones missed by target sequencing estimated as the difference between the number of clones predicted by our model and the number of clones detected from target sequencing samples collected from 1,923 donors.

We applied this ABC scheme independently to the two distinct datasets. The first comprised ∼ 700 variant trajectories from ∼ 400 donors (Fabre et al. [15]). We compared these with simulations of Equation (1) using two summary statistics: the distribution of clone sizes across all donors and the distribution of logistic fitness coefficients *s*_log_ (Figure 3B-C). The second consisted of single-cell expanded HSC colonies from 8 individuals aged 0-81 (Mitchell et al. [16]). For these, we generated longitudinal samples of our agent-based simulations to compare the cumulative SFS (Figure 3D, Figure SI 9A-H) and the number of detectable clones above 1% frequency (Figure 3E, Figure SI 9I). See SI 5.4 for details. Best-fit simulations capture the individual stochasticity of clonal expansions that manifest as subclonal peaks in the higher frequency domain of the SFS (Figure SI 10). They furthermore accurately predict how the entropy of the HSC pool changes with age, revealing a pronounced deviation from the neutral expectation later in life (Figure 3J).

The posterior distributions for the competition parameters (*η, σ*, and *µ*) converge across both datasets and deviate significantly from imposed uninformed priors in both ABC inference schemes (Figure 3F-H, and Figures SI 2, SI 11). We report mode and 90% credibility intervals for best parameter values. From the inference on variant trajectories (Figure 3B-C) we find for the mean and the standard deviation of the fitness distribution 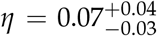 per year and 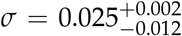 (Figure 3F-G). For the arrival rate of advantageous variants in the stem cell population we find 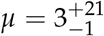 per year (Figure 3H). The inference on the SFS (Figure 3D-E) estimates 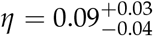 per year, 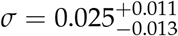 (Figure 3F-G), and an arrival rate of advantageous variants of 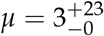 per year (Figure 3H). The posterior parameter distributions for the HSC population size *N* and the HSC wild type inter-division time *τ* converge less sharply (Figures SI 2, SI 3, SI 11). This is in accordance with our validation of the ABC framework on synthetic data (SI 3.1, Figure SI 4B), and implies that the competition model is compatible with a wider range for these parameters.

The inferred innate fitness distribution peaks at a positive non-zero value (Figure 3I), implying that clones with very low innate fitness (*s* ≈ 0) are rare. To ensure this result is not due to limited data resolution – since low fitness clones are less likely to expand to observable frequencies – we performed ABC inferences on simulated data with an imposed exponential innate fitness distribution measured at similarly limited resolution. The ABC successfully identified the correct innate fitness distribution despite the lowest fitness variants not being directly observed (Figure SI 4A). This demonstrates that even low fitness variants convey a measurable effect on the system dynamics through compounding interactions with other clones. It is worth noting that – due to the nature of the data – the inferred distribution (Figure 3I) describes the prevalence of advantageous variants that have not yet transformed into aggressive cancers, and thus potentially does not describe the fittest variants that likely occur at high *s*. This agrees with recent estimates of AML subclonal driver mutations with estimated fitnesses between 0.4 to 0.8 [27], and estimates of the most common mosaic chromosomal aberrations in blood that aggregate at the upper bound of 0.1 to 0.2 per year of the fitness distribution [28].

The parameter inference finds an arrival rate of fit clones on the order of 2 to 20 per year, suggesting that by age 50 an individual may have harboured up to a thousand clones under positive selection. Though this may appear high, the fraction of fitness-conferring mutations is still low overall. In a population of 10^5^ HSCs with a total rate of 20 mutations per year per stem cell [5], this estimate results in 1 to 10 in 10^6^ mutations conferring a selective advantage. Furthermore, the estimated fitness distribution has its average at 7-9% per year. Thus the combined effects of low fitness and competition prevent most clones from escaping extinction and reaching detectable sizes.

Since our model provides an expectation for the number of expanded clones conditional on age, we sought to compare this quantity against larger datasets. We applied our framework to 1,942 variant trajectories, integrating the previously analysed trajectory data [15] with two additional public datasets [17, 21] (see SI 6). The number of expanded clones in the data suggests that targeted sequencing current panels of haematopoietic driver mutations likely misses as many as 10 expanded clones exceeding a frequency of 1% by the age of 75 (Figure 3K). This is corroborated by the still ongoing discovery of novel CH drivers [29] and the high prevalence of ‘driverless’ CH [16]. Some of these undetected clones may derive from non-targeted genic regions, non-coding regions, structural variants, or epigenetic modifications.

### Innate fitness and arrival time estimation of 1,942 variant trajectories

We next explored whether innate fitnesses of clones are identifiable in a dynamic HSC fitness landscape. Complications arise because trajectories depend on both innate fitness and competitive interactions with other clones, many of which may be undetected. The parametrised clonal competition model, however, predicts the distribution of lifetime behaviours of a clone arising at a particular age and fitness. This enables inference of the innate fitness most likely to result in a given trajectory. We first tested this approach against simulated data. We embedded 1000 clones with predetermined innate fitness *s* and arrival time *t*_0_ in independent realisations of the SDE model, to ensure the ensemble encompasses the inherent stochasticity of clonal competition (Figure 4A). This resulted in a set of clone trajectories with known initial conditions. We compared this *in silico* dataset to 150,000 simulated trajectories with uniform randomly drawn innate fitnesses and arrival times. Taking the maxima of the resulting marginalized posteriors (Figure 4B), we find the distribution of estimates to centre around imposed values, with increasing variance for higher fitness and earlier arrival times (Figure 4C).

**Figure 4.**
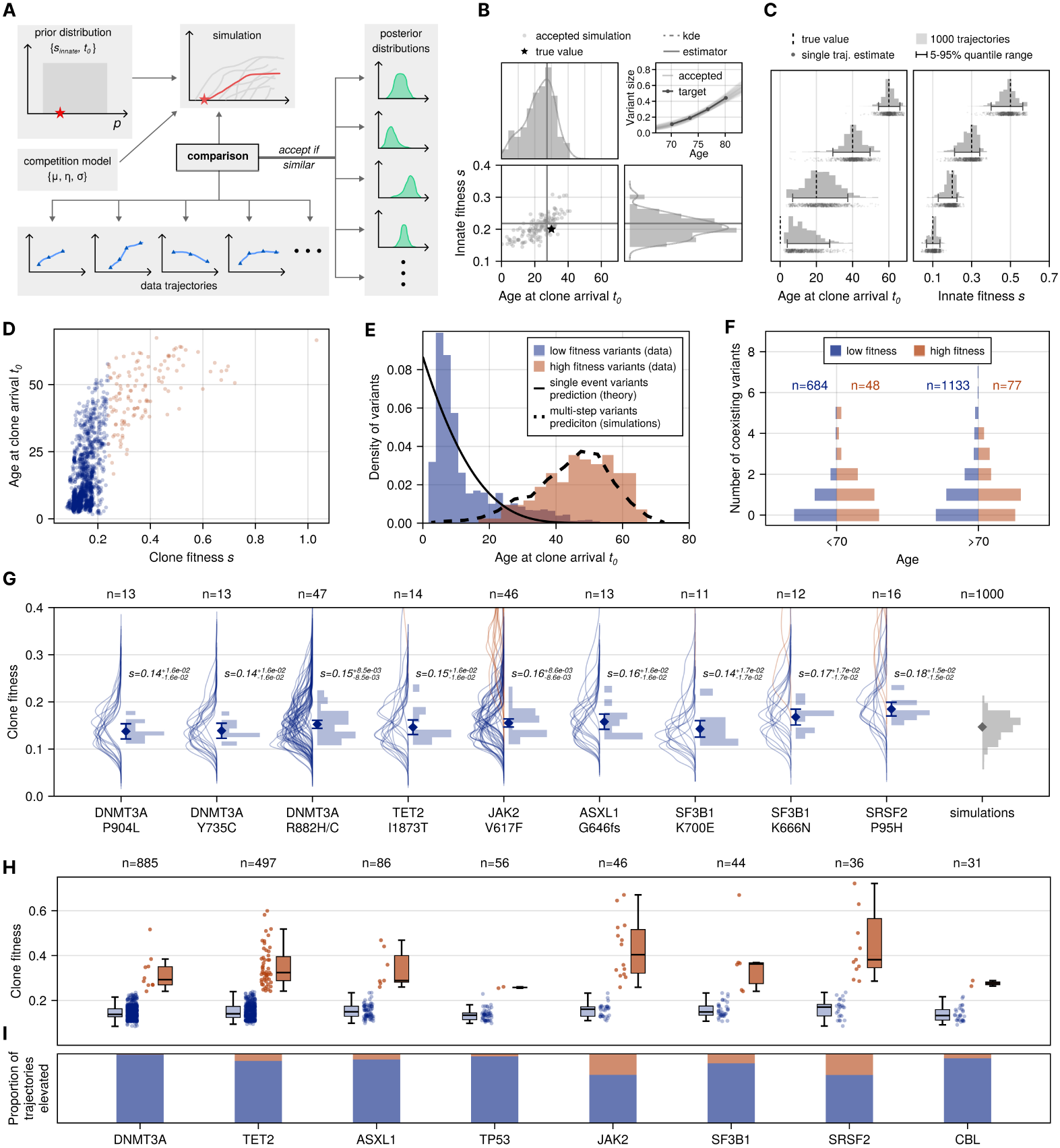
Individual variant and genic innate fitness estimates and timing. **A)** Schematic of the ABC framework to infer arrival time *t*_0_ and innate fitness *s* of individual variants. **B)** Example outcome of the ABC for a single simulated variant trajectory. **C)** Distribution of ABC estimates on simulated data with fixed values (dashed vertical lines, 1000 trajectories per value). **D)** Estimates of arrival time *t*_0_ and fitness *s* of 1,942 variant trajectories from three datasets (Fabre, Roberston, Van Zeventer). Trajectories are segregated in either low (*s* < 0.24, blue circles) or high (*s* > 0.24, red circles) fitness. **E)** Distribution of arrival times of both low (blue bars) and high (red) fitness variants, compared to predictions from theory (solid line) and simulations (dashed line) for single and multi-step clonal evolution. Te distributions are normalized independently for better visualisation. **F)** Distribution of the number of observed variants that coexist with the target variant, for low (blue left facing bars) and high (red right facing bars) fitness, in individuals under (left) and over (right) 70 years of age. **G)** ABC-derived posterior distributions (left facing solid lines) for innate fitness *s* of most common identical variants across multiple individuals (number shown above). Right facing histograms show distributions of innate fitness estimates (maxima of posteriors). Rangebars denote twice the standard error above and below the mean (diamonds). Estimates of *in silico* variants (far right, grey) from 1000 simulated trajectories with random arrival times *t*_0_ ∈ [0, 60] and innate fitness *s* = 0.15. **H)** Estimated innate fitnesses stratified by most commonly mutated genes. **I)** Per gene proportions of low (blue) and high (red) fitness drivers.

We applied this framework to the 1,942 variant trajectories in the aggregated dataset [15, 17, 21] (SI 6). For all trajectories the marginalized posterior distributions converged in both parameters. The majority (1,817 out of 1,942) had innate fitness estimates between 0.08 and 0.24, placing them above the mean of the fitness distribution, in line with the expectation that only sufficiently fit clones reach detection (Figure 1G). Only a small subset of variant trajectories (125 out of 1942) presented with higher fitnesses (> 0.24). Notably, these tended to arise much later in life, whereas lower fitness trajectories scattered across all ages (Figure 4D, E). This observation is remarkably well quantified by a multi-step model of clonal evolution. The arrival time distribution of the lower fitness variants agrees with the analytical expectation of single events occurring with constant rate throughout life. While the arrival times of high fitness variants match simulated expectations of a multi-step process (Figure 4E, SI 8). Such a multi-step model of clonal evolution is further supported by clonal co-occurrence patterns. In donors under the age of 70 the high fitness trajectories had a higher likelihood of being detected alongside at least one other driver (61% of highest fitness trajectories versus 42% in other drivers, *p* = 0.012 Mann-Whitney *U* test), and had a higher average number of co-occurring drivers (1.13 versus 0.62, *p* = 0.004 Mann-Whitney *U* test, Figure 4F). These statistical differences vanish at older ages (> 70), likely as individuals harbour increasingly more detectable clones masking correlations of co-occurrence patterns. Taken together, this suggests that the lower fitness trajectories (< 0.24) are mostly derived from single events, and their estimated fitnesses may be identified as the innate fitness of their driver. In contrast, for higher fitness variants (> 0.24), multiple steps have elevated their fitness above the driver’s solitary innate value.

### Variant innate fitness variation and elevated fitness prevalence

Identical genic variants present with largely varying growth rates across individuals and age [15, 21]. We have shown that competition-driven changes in the HSC fitness landscape promote such disparity. But how much of the observed heterogeneity is explained by competition? To address this, we examined identically mutated sites across multiple donors. The variants most commonly occurring (n≥11) among the 1,942 trajectories comprised three mutations in DNMT3A (n=47, n=13, n=13), two mutations in SF3B1 (n=12, n=11) and single mutations in JAK2 (n=46), SRSF2 (n=16), TET2 (n=14), and ASXL1 (n=13). We obtained independent distributions of fitnesses (Figure 4G) for all distinct variants, e.g. for the DNMT3A.R882H/C mutation this gives us 47 posterior fitness distributions. From these we took the mean across donors – excluding multi-step trajectories (*s* > 0.24) – as the variants’ innate fitness. The estimated innate fitnesses ranged from 0.14 to 0.18, with standard errors between 0.008 and 0.016 (Figure 4G, Table 2). For a single variant, the variation between individual trajectory estimates is in part caused by uncertainty from the inference method due to stochasticity of the competition model itself (SI 7). Quantifying this allows us to infer how much variation would remain in the absence of competition. To this end we performed the same innate fitness inference on 1000 simulated trajectories with random arrival times and fixed innate fitness *s*=0.15, resulting in a distribution with mean inferred fitness 0.148 and standard deviation 0.026 (Figure 4G). In comparison, the average standard deviation across all nine investigated variants was 0.029, and the two most abundant variants in the data – JAK2.V617F and DNTM3A.r882H/C (46 and 47 donors as opposed to ≤ 16 donors for others) – had standard deviations of 0.029 and 0.028 respectively. This implies about 80% of the variance on single variant estimates can be explained by competition (see SI 7.2).

Surprisingly, none of the combined 73 trajectories of the most common DNMT3A variants had elevated fitness (Figure 4G). In contrast, for JAK2.pV617F, SRSF2.p95H and TET2.1873T we observe trajectories of elevated fitness despite small sample sizes. The same trend remains if we compare variant fitness across genes. Only 10 out of 885 DNMT3A variants have elevated fitness. In contrast, 51 out of 497 TET2 variants and 11 out of 36 SRSF2 variants have elevated fitnesses. Among the nine most commonly mutated genes, mean elevated fitness ranges from 0.26 to 0.49, with noticeable variation between genes. The ranges are consistent with recent retrospective fitness inferences of preleukemic clones with two or three accumulated driver mutations that would later transform into AML [30]. It is encouraging that these high-risk multi-step events seem identifiable prior to progression, raising hopes for risk stratification of CH trajectories across individuals and age.

## Discussion

The clonal competition model ultimately arises from the notion that the HSC population is strictly regulated to approximately maintain its size in adults. This is perhaps most strongly illustrated by the bone marrow’s ability to repopulate its HSC pool following transplant [31]. It is likely that particular non-malignant or pre-malignant variant combinations allow partial escape from such regulation. However, a model of unconstrained exponential growth for all advantageous variants would result in a 5- to 100-fold increase of the stem cell population late in life (Figure SI 14), for which there is currently little evidence in human haematopoiesis.

The converging inferences of the CH fitness distribution and clone arrival times in two different *in vivo* datasets suggests that clonal competition naturally explains complex and contrasting observations in ageing haematopoiesis. It also suggests that CH and age-related expansions are ubiquitous in the elderly. Our model explains the sudden transition in the number of detectable clones between the ages 50-60 [13, 16], and provides sharp bounds on the possible number of detectable clones in the elderly. Above the age of 70, between 5 and 20 clones should be visible at a resolution of 1% VAF (Figure 3E). It furthermore correctly predicts the non-monotonic change of entropy within the HSC population with age (Figure 3J). Interestingly, entropy initially increases analogous to a neutral process – which sets the maximal possible rate – and only deviates by the age of 63 as clonal expansions become more prevalent. Noteworthy, simulations suggest a very small variance of the entropy during adulthood, possibly providing a method to detect abnormal HSC dynamics early.

The inference of innate fitness from single trajectories forms an important step towards better understanding the behaviour of CH drivers. As opposed to other methods of fitness estimation – such as function fitting or growth rate measurement – our approach accounts for the stochastic noise introduced by individual-specific clonal competition. From these estimates it is striking that 80% of the variance (approximately 90% of the standard deviation) observed on identical variants can be attributed to competition.

The clear distinction in arrival times between low and high fitness variants highlights the role of accumulating driver events along the path of neoplastic transformation, and the importance of better understanding this process. For example, the considerable under-representation of DNMT3A variants among elevated fitness trajectories is a surprising observation that warrants further investigation. DNMT3A.R882 has high innate fitness and plays a prominent role in transformation to AML. Yet, despite its dominant prevalence in CH, it appears to co-occur less frequently with many other AML driver mutations [32]. This might speculatively point to a rugged fitness landscape of DNMT3A and co-occurring variants – with few viable variant combinations – implying a slow transition rate and higher prevalence of single-step DNMT3A variants in CH. Conversely, the absence of elevated-fitness DNMT3A trajectories may also point to a survivorship bias, where rare jackpot events lead to fast progression. It also remains an open question if all DNMT3A.R882 variants are equally likely to transition, or if there are differences in the patterns of hypomethylation that prime some over others. Answers may come from evolutionary models of clonal competition combined with single cell analysis [33]. We expect that with higher resolution additional effects such as age-related HSC deregulation, immune interactions, inflammation, epistasis, and frequency-dependence may eventually become identifiable as well.

Lastly, it has been speculated that clonal competition may act as a barrier to some malignant expansions, since widespread competition amongst a large population of HSCs limits growth of individual clones [34]. This work provides some credence to the idea, however the strength of this effect is not clear. Furthermore, its exact manifestation may be highly tissue-specific, and thus the extent to which it might impede the transition from healthy tissue to malignancies remains an open question.

## Acknowledgments

N.V.MP. and F.T. acknowledge support by the Barts Charity. B.W. is supported by a Barts Charity Lectureship (grant no. MGU045) and a UKRI Future Leaders Fellowship (grant no. MR/V02342X/1). We are grateful to David Dingli, Andrea Sottoriva, Alexander Stein, Jamie Blundell, Eric Latorre Crespo, Linus Schumacher and anonymous reviewers for discussions and comments at different stages of this work.

## Code availability

The HSC dynamics are implemented in the Rust programming language and the code is available at https://github.com/fraterenz/hsc. The numerical implementation of the SDE system in Equation (S18) was written in the Julia programming language and is available at https://github.com/natevmp/competitive-selection. The Python code to recreate the figures is hosted at https://github.com/fraterenz/hsc-draft.

## Supplemental Information

## SI 1 Maximum likelihood estimation of logistic trajectories

The logistic function – obtained as the solution to the differential equation d*x*(*t*)/d*t* = *s*_log_ *x*(*t*)[1 - *x*(*t*)] – describes the expected growth curve of a mutant under selection in a constant-size population with fixed background fitness [1]. Because the observed variant trajectories are sigmoidal, it can be used to approximate a trajectory in a short time window. Integrating the differential, the clone size is given by

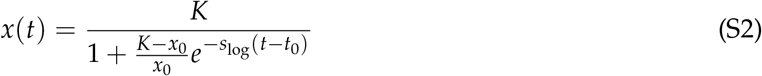

where *x*_0_ is the clone’s initial size at time *t*_0_ and *K* is the carrying capacity. Importantly this expression holds for both positively and negatively selection mutants, since in case of the latter flipping the sign of *s*_log_ is akin to flipping the sign of *t*, and at any point in time both subpopulations must have their sizes sum to *K*. If a trajectory is well-described by Equation (S2), the rate parameter *s*_log_ functions as a measure for the selection strength acting on the clone in the observed time window.

We fit all variant size trajectories reported by Fabre et al. [2] to Equation (S2) using a maximum likelihood estimation as follows. We assumed each measurement of a variant allele’s frequency 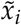 to be a binomial sampling, where the success probability is given by the (unknown) true frequency *x*_*i*_ and the number of trials is given by the coverage *V* of that site. Thus for any proposed trajectory *y*(*t* | *β*) (where *β* is the set of parameters to fit), we could compute the probability of obtaining the observed sequence of measurements 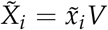 at times *t*_*i*_ as

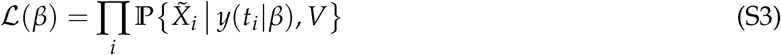

where

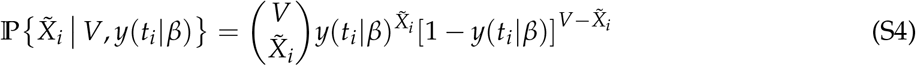

which is the binomial probability of sampling successfully 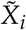 times out of *V* trials with success probability *y*(*t*_*i*_ | *β*). Equation (S3) is known as the likelihood function, and is maximized with respect to *β* to obtain the best fit. For each trajectory we maximized three different likelihood functions: one assuming the underlying trajectory is increasing, another assuming it is decreasing, and finally one assuming it is constant (*y*(*t*) = *x*_0_). In the case of an increasing trajectory on a non-sex chromosome we took *K* = 0.5 and *x*_0_ = 1/2*N* as fixed parameters and – assuming *s*_*log*_ > 0 – maximized with respect to *β* = { *t*_0_, *s*_*log*_ }. For a decreasing trajectory we took *β* = {*t*_0_, *x*_0_, *s*_*log*_ } (*x*_0_ is then the size at which the clone became negatively selected) and instead assumed *s*_*log*_ < 0. Finally for the saturating trajectory *s*_*log*_ = 0 and the single free parameter *β* = { *x*_0_ } (the fixed true clone frequency) was fit. The trajectory with the highest maximum likelihood was then taken as the best fit.

## SI 2 Clonal competition as a multi-type capacity-limited birth-death process

We model the haematopoietic stem cell compartment as a collection of cells, each carrying a set of genetic mutations. These cells individually undergo divisions and death as independent Poisson processes in time, where in case of the former the cell is replaced by two daughter cells inheriting the set of mutations from the parent, and in case of the latter the cell is removed from the population. We consider first only fitness-granting mutations which alter the birth rate of a cell. We take these to occur as a Poisson process in time, and assume each new mutation is distinct from those already existing, a supposition typically referred to as the *infinite sites model*. Denoting the set of all fitness carrying mutations as ℐ, a cell of type *i* ∈ ℐ as 𝒳_*i*_, and the birth, death, and mutation rates as *β, δ*, and *µ* respectively, the model is summarized through the following reactions:

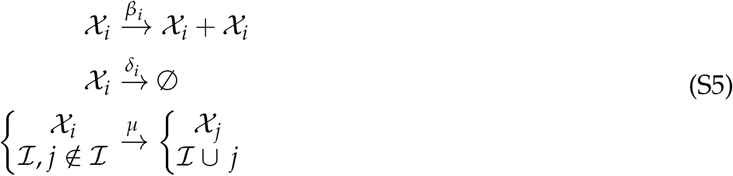

An important constraint is that the total stem cell population is expected to remain at a constant size throughout most of the human lifespan [3, 4]. We take this to occur by enforcing an equality of the total population-level birth and death rates:

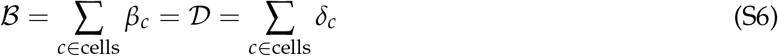

with *β*_*c*_ and *δ*_*c*_ the birth and death rates of cell *c*. If all cells are of the same wild type – having identical birth and death rates – this is trivially realised by setting *β*_*c*_ = *δ*_*d*_ = *β* = *δ*, ∀*c, d*. This amounts to a completely neutral population, where ℬ = *Nβ* = 𝒟 = *Nδ* for a population of size *N*. If there are *K* types the population-level rates can instead be written as

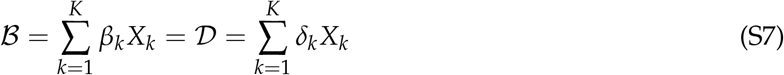

where *X*_*k*_ denotes the number of cells of type *k*, and all cells of that type share the birth and death rates *β*_*k*_ and *δ*_*k*_. We define selection acting on a type *k* as an altered birth rate with respect to the wild type through a fitness parameter *s*_*k*_, and a fixed death rate across all types ∀*k* : *δ*_*k*_ = 𝒟/*N*:

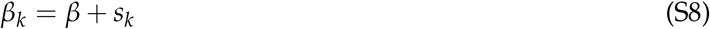

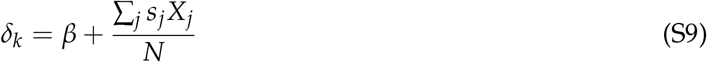

### SI 2.1 The SDE system for clonal competition

#### SI 2.1.1 Deriving the SDE system

From the rates (S8) and (S9) a master equation can be constructed for the probability distribution of the size of type *k*, where we use the shorthand notation 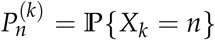 :

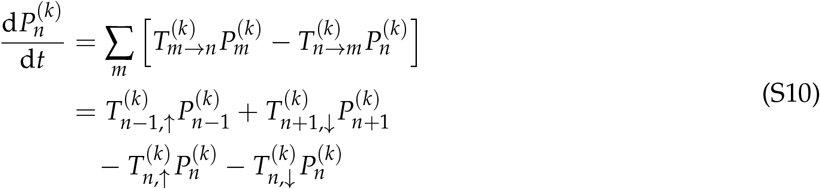

This expression contains the transition rates 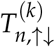 for the size of clone *k* to increase or decrease by 1:

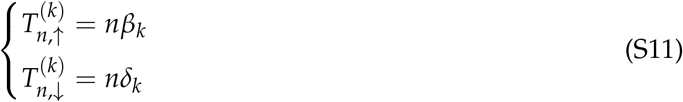

Assuming that *N* is large, meaning that there are many possible values of *X*_*k*_/*N* between 0 and 1, we construct an analogous stochastic process where the size is given by a continuous variable *x*_*k*_ ∈ [0, 1] ⊂ ℝ, representing the frequency of clone *k* in the population. One convenient way to do this (see for example [5]) is by writing the master equation as a Kramers-Moyal expansion. First, noting that the transition rates here describe displacements of distance Δ*x* = 1/*N*, i.e. 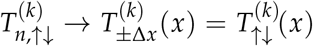, we rewrite (S10) in terms of a Taylor expansion of the continuous parameter *x*:

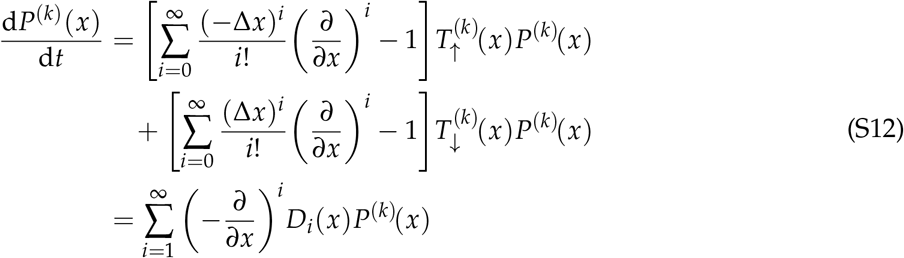

where the final equality immediately describes the Kramers-Moyal expansion with coefficients

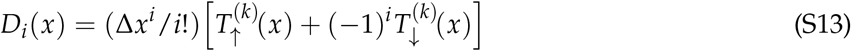

Using our assumption of small steps Δ*x* = 1/*N* we truncate after the second term, to obtain a Fokker-Planck equation with coefficients

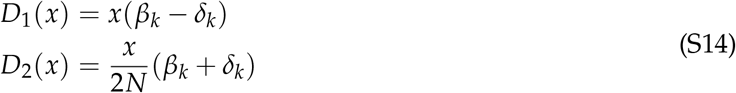

These can be used to construct an equivalent stochastic differential equation

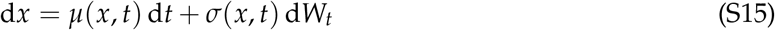

where in the Itô formulation the drift and diffusion coefficients take the form

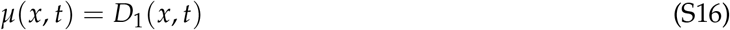

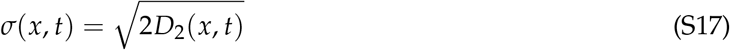

Thus, plugging in (S8) and (S9), we have

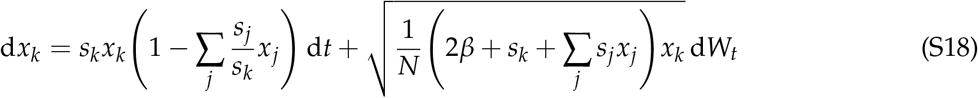

which describes a set of coupled stochastic differential equations for *k* = 1, 2, …, *K*.

#### SI 2.1.2 Numerical simulations of the SDE system

With Equation (S18) describing the time-evolution of existing variants clones, all that remains to obtain the model described by Equation (S5) is the introduction of newly arriving clones in time. It suffices to take the total number of equations *K* to be a Poisson process in time, and have newly introduced clones initiate at size 1/*N*:

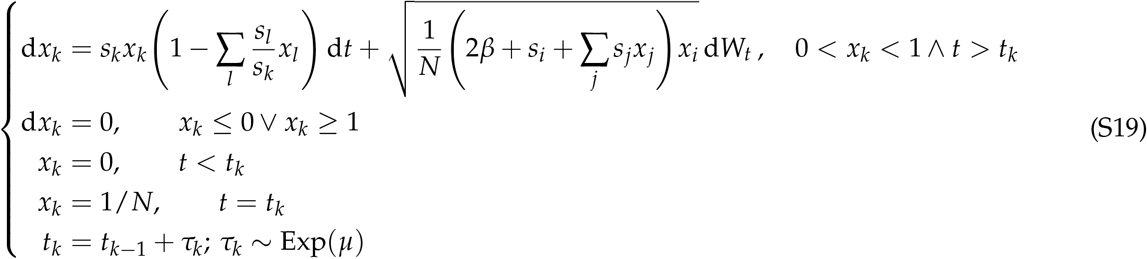

Upon introduction, a clone *k* has its fitness *s*_*k*_ drawn from a gamma distribution *s*_*k*_ ∼ Gamma 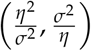 with mean *η* and variance *σ*^2^. Numerical solution trajectories of this system were calculated using the Julia computing language and the package ‘DifferentialEquations.jl’ [6–8].

### SI 2.2 Unconstrained growth modeling

An alternative proposed model for CH [9, 10] is one in which there are no constraints on the total number of HSCs, effectively allowing all fit mutant clones to grow exponentially without competition. Because size changes are measured in terms of allele frequencies, growth trajectories would still appear sigmoidal upon measurement [10], suggesting such a model may predict similar empirical observations. However, an important difference is that in such a scenario the total HSC population must growth exponentially along with the expansion of its inhabiting mutant clones. As there currently exists little evidence for such behavior, we investigated to what extent the HSC pool would be expected to expand in a human lifetime under such a model. Constructing a birth-death process and moving to a diffusion picture in a manner analogous to SI 2.1, it is straightforward to show that the absolute number of cells *X*_*k*_ in a clone obeys the SDE

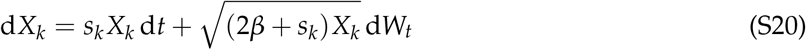

with *β* again the wild type division rate. The observable frequency of a clone can then be found as *X*_*k*_/*N* = *X*_*k*_/ ∑_*j*_ *X*_*j*_, where *N* = ∑_*j*_ *X*_*j*_ includes *X*_0_, the fixed wild type population size. Using the optimal parameters obtained from ABC inferences (Figure 3H-J) we measured the distribution of total population sizes over time across 1000 simulations (Figure SI 14). Taking the initial mature HSC pool to house 100’000 cells, we find a large variation of increases in this size across 1000 simulations. Even in the scenario of lowest clone arrival rate (*µ* ≈ 3 per year) the median population increase is 5-fold that of the initial size, with some simulations reaching 100-1000 fold increases. Such drastic changes to the composition of the bone marrow would likely effect other aspects haematopoiesis, a notion which warrants further empirical investigation.

## SI 3 Bayesian inference on clonal trajectories to estimate parameter values

The Bayesian inference on the clone trajectories (Fabre et al. [2]) was performed in the following manner. First 40’000 parameter combinations {*P*_*i*_ }– hereafter referred to as particles – were drawn randomly from uniform priors for each of the *n* parameters to be investigated. Next, for each particle a simulated dataset was generated from the SDE system, from which the variant size-distribution and logistic fitness distribution were measured and compared to those of the donor data using a *χ*^2^ metric. Finally a threshold was selected such that only the 1% best performing particles (i.e. having the highest similarity with the Fabre data according to the metric) were retained, resulting in a *n*-dimensional posterior distribution of the parameter space (Figure SI 2). Sensitivity of the posteriors was tested by varying the number of particles accepted, at 0.1%, 0.5%, and 5% (Figure SI 5).

To construct the simulated dataset for a single particle, 300 realisations of the SDE system were performed, representing a large cohort of virtual donors. Each run was measured at time *T*, and of the variants exceeding a detection threshold *x*_*min*_ = 0.002, only up to three were chosen for tracking, to obtain a similar number of total variants as well as variants per donor in the data. Because trajectories in the dataset were captured through targeted sequencing of known driver genes, it is possible that the observed variants would be biased towards higher innate fitness compared to a randomly selected set of drivers, since knowledge of a driver may correlate with its fitness. To account for this we introduced another parameter *q* describing the probability of an observed driver possessing the highest innate fitness in the system. In concrete terms, after ranking all variants in a simulated individual by their innate fitness, each of the observed variants has a probability *q* of being chosen in ranked order, and a probability 1 − *q* of being selected randomly. Thus for *q*=1, observed variants are maximally biased towards high innate fitness – i.e. sampled in order of innate fitness – whereas for *q*=0 variant observation is entirely random. From the pooled set of selected variants across all simulated individuals, the size distribution was constructed by binning variant sizes at the time of measurement into 25 bins covering the full VAF range [*x*_*min*_/2, 0.5] (the size of a clone is twice its variant allele frequency). For the fitness distribution, each sampled variant was measured at the four equidistant time points spanning 10 years centered around T: {*T* − 5, *T* − 1.66, *T* + 1.66, *T* + 1.66 }. The maximum likelihood fitting described in SI 1 was then performed to obtain all variants’ logistic fitness coefficients *s*_log_. which were partitioned into 50 equidistant bins constituting the fitness distribution. Because both the size- and logistic fitness distributions are expected to change with the age of observed individuals, the donor dataset was partitioned into age groups covering 6 year spans, and the comparison with simulations was performed with *T* at the center of the respective age group, for example variants measured between 70-76 years of age were compared with simulations measured at *T* = 73.

### SI 3.1 Testing the ABC convergence with simulated data

To test the ability of the ABC framework to accurately identify unknown parameter values, we performed the inference on 1000 simulated datasets with known parameters. As inferred values we took the modes of the posterior distributions. Taking the Euclidean distance between inferred and true parameter value as the error, we then measured the distribution of errors across all inferences for each parameter (Figure SI 4B). For all parameters the error distributions have modes near 0 and mean errors less than 1 fold-change.

### SI 4 Accumulation of neutral mutations in the multi-type birth-death process

To study site frequency spectra (SFS) we must include the accumulation of neutral mutations in individual haematopoietic stem cells, as these are expected to form the majority of the cell mutational burden. Current evidence suggests mutations can occur both during cell division due to replication errors, as well as in between divisions due to persistent background stresses on the genome (e.g. radiation, oxidation, etc.) and imperfect DNA repair [11]. We must distinguish between these two forms of mutation as they lead to different distributions of the mutational burden in single cells [12]. The number of division-associated mutations in a cell scales with the number of divisions its parental lineage has undergone, whereas the number of background-associated divisions scales directly with time. We take division-associated mutations to be acquired as a Poisson distributed number during each division, and background mutations as a Poisson process in time. To this end we expand upon the events described in Equation (S5). Denoting 𝒳_*k*,{*u*}_ a cell of type *k* with set of mutations {*u*}, and *U* the set of all existing mutations, division-associated mutations are introduced during birth events

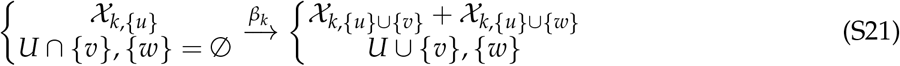

and background mutations establish the additional event

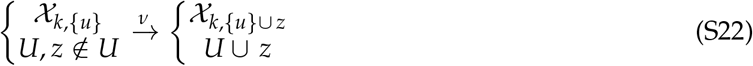

### SI 4.1 Ontogenic growth of the HSC population

Because population expansion drives neutral mutations to high frequency, the shape of the SFS reflects changes in a population’s size [13]. These changes can occur gradually, such that SFS measured from blood in adult humans still contain signatures of the early development of the HSC pool. In order for a model to accurately capture the changing SFS in an individual, it must therefore also incorporate early ontogenic growth [12]. To this end we include an exponential growth phase preceding the previously-described constant population dynamics, starting from a single cell and lasting until the adult size of the HSC pool *N* is reached, which we take as the time of birth (0 years old). We take this phase to occur as a pure-birth process without cell death.

### SI 4.2 Agent-based model with inclusion of neutral mutations

We implemented the clonal competition model with accumulation of neutral mutations as a Moran model [14] via the stochastic simulation algorithm (SSA, also referred to as the Gillespie algorithm [15]). The SSA generates stochastic trajectories of the model by iteratively simulating the events described in Equations (S5), (S21), and (S22) in a system of cells. Both the timing between events and the events themselves are drawn randomly, so that each simulation is a unique realisation of the underlying stochastic process. In our model we assume a *pure-birth process* for the ontogenic growth phase: each event describes a cell division and results in the appearance of a new cell in the system. In the constant-population phase we assume *Moran* dynamics: each event consisting of a birth and a simultaneously occurring death, which enforces an equality of the total birth and death rates and thus keeps the population size constant over time. All events are assumed to occur as Poisson processes in time, i.e. having exponentially distributed waiting times. This means that following an event at time *t*, from Equation (S21) the time until the next event is given by

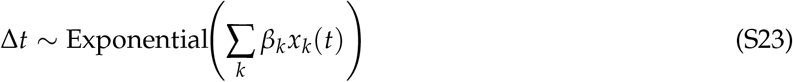

with *x*_*k*_ denoting the size of clone *k* and *β*_*k*_ its birth rate. The clone to which the proliferating cell belongs is then drawn randomly from the weighted set of existing clones, with the likelihood of clone *k* being selected given by *x*_*k*_*s*_*k*_/ ∑_*l*_ *x*_*l*_*s*_*l*_. Repeating this sampling procedure generates a sequence of proliferating cells and inter-division times, representing a random realisation of the process described by Equation (S10) without requiring an analytical solution for 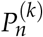.

We implemented the SSA in the Rust programming language [16, 17] as an agent-based model (ABM). Each agent represents a single HSC, which carries a set of mutations {*u*} and belongs to a clone *k*.

The proprieties of the agents define the state of the system at certain time, and we are interested in understanding the changes over time of the state of the system. Upon proliferation, a random cell belonging to the *k*-th clone is selected and assigned *m*_*b*_ ∼ Pois(*λ*_*b*_ Δ*t*) background neutral mutations, where 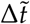 is the difference between the current time and the last time the cell divided. Then a Bernoulli trial with probability of success *µ*Δ*t*/*N* is performed to determine whether to allocate the cell to a new empty clone. The dependence on the stochastic inter-division time decouples the occurrence of fit clones from proliferation events, ensuring that our framework is agnostic to the type of event that generates new clones: they do not acquire a proliferative advantage exclusively via division-associated genetic mutations, but also due to background mutations, epigenetic alterations, or other environmental factors. When a novel clone is created its fitness *s*_*k*_ is drawn from a Gamma distribution. This quantifies how fast cells in the clone proliferate according to Equation (S8). Finally, the dividing cell is duplicated and each daughter cell is assigned a set of random mutations *m*_*d*_ ∼ Pois(*µ*_*d*_). If the system has passed the growth phase, a random cell drawn uniformly across all clones is removed from the population. Since we generate clock-like mutational events with the background mutations *m*_*b*_, we ignore asymmetric divisions for simplicity, which would only increase the mutational burden. Repeating these steps many times allowed us to simulate both clonal dynamics and the accumulation of neutral mutations that act as lineage markers in individual cells.

## SI 5 The single-cell HSC dynamics

Having extended the model to incorporate distinct mutations in individual cells, we aimed to understand the dynamics of clonal haematopoiesis (CH) from single-cell resolution data published in Mitchell et al. [4]. We show that using different summary statistics – namely the single-cell mutational burden, the site frequency spectrum and the number of expanded clones – a clonal competition model can quantify underlying evolutionary parameters of CH and recapitulate the genetic heterogeneity observed in the data.

### SI 5.1 The dataset of whole genome sequencing of HSC colonies

The HSC whole genome sequencing data published by Mitchell and colleagues [4] consist of samples collected from either the bone marrow, the peripheral or cord blood of 9 healthy donors aged 0 to 81. Cells were sorted and colonies were grown from individual haematopoietic stem cells or multipotent progenitors (HSCs/MPPs) and from haematopoietic progenitor cells (HPCs). For this study we assumed that both HPCs and HSCs/MPPs are subject to the HSC dynamics as differentiated cells in blood show similar mutation loads to HSCs [11]. Colonies were whole-genome sequenced providing high coverage data at single-cell resolution: only clonal colonies were kept for further analysis resulting in 3’579 sequenced colonies, of which 3’361 were grown from HSCs/MPPs and 218 from HPCs. We downloaded the genotype matrices of 9 individuals (3’046 cells) from Mendeley https://data.mendeley.com/datasets/np54zjkvxr/2 and used them to compute the single-cell mutational burden (that this the number of distinct mutations in individual cells) and the site frequency spectrum. We also stored the number of expanded clones per donor, defined as the number of postnatal ancestral lineages whose descendants contributed more than 1% of colonies at the time of sampling. Note that the information about the clones is extracted from the phylogenies reconstructed by [4] from the single-cell mutations.

### SI 5.2 The HSC site frequency spectrum dynamics

We quantified HSC genetic heterogeneity using the site frequency spectrum (SFS) – the distribution of prevalences of all somatic variants within a sample. We analysed the SFS of the donors to assess whether a clonal competition model can recapitulate the genetic heterogeneity observed in the data.

The SFS were computed from the single-nucleotide variants and short indels present in the individual cells of the 9 donors (216-451 cells per donor, 362 cells on average) stored as the genotype matrices. For each donor, we calculated the frequency of cells carrying each individual mutation, and then counted all mutations having the same frequencies to obtain the variant counts. We then computed the variant frequencies by normalising the variant counts.

Figure SI 6 shows the SFS of the donors (orange crosses) along with individual simulations and averages obtained from the ABM model (grey dots and lines). The theoretical expectations of the SFS for expanding populations under neutral dynamics (solid black lines), which follow a 1/ *f* ^2^ power-law distribution [13, 18], were sampled to match the sample size of each donor. The theoretical expectations of the SFS for a fixed-size population (dotted black lines) were computed by simulating a birth-death process without selection starting from a first growing phase followed by a fixed-size population phase [12] and were sampled to match the sample size of each donor.

### SI 5.3 Model parameterisation with the mutational burden

We aimed to quantify the parameters of the ABM clonal competition model (Table 1) from the single-cell data described above. Four parameters were estimated from a Bayesian model (see SI 5.4), while the others were either fixed from the literature or parameterised using additional information from the data. From current knowledge we set the rate of neutral mutations upon division *µ*_*d*_ to 1.14 [3, 19], and fixed the number of stem cells *N* to 10^5^ cells, based on recent estimates [3, 4, 9, 12] and the fact that the ABC inference performed on clone trajectories displayed limited sensitivity to this parameter within a wide range (Figure SI 3). Finally, we parameterised the rate of accumulation of neutral background mutations using single-cell mutational burden information from the data (Figure SI 7A). To accurately capture the dynamics of the burden observed in the data, we assumed different values of this parameter before and after birth, denoted 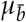 and *µ*_*b*_ respectively.

To compute the prenatal background mutation rate 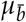, we took mutations to occur as following: an initial growth phase, lasting *t*_growth_, has mutations occurring through both divisions and background processes until the maximal population size *N* is reached; then for the remaining time until birth, that is *t*_birth_ − *t*_growth_, only background mutations are added. Having the growth process complete before birth (*t*_growth_ < *t*_birth_) allowed for better agreement of the SFS between simulations and data from newborns. We chose *t*_growth_ such that simulations match the SFS of the neonates in the data, which we found to be 5 months (Figure SI 6A-B). The value of *µ* is then obtained from

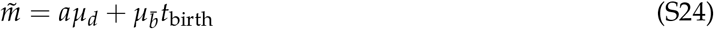

where 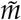 is the expected burden distribution upon birth, *t*_birth_ is the total time elapsed before birth (9 months), and *a* = 2 log ((*N* + 1)/2) the expected number of divisions per lineage once the population has reached size *N* [12]. Taking for 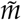 the mean of the burden distributions measured from the two newborns in the data (Figure SI 7A) resulted in 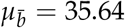.

To compute the postnatal background mutation rate *µ*_*b*_, we regressed the means of the burden distributions for all individuals with no detected clones (i.e. individuals with 0, 29 and 48 years). We found an increase in the average burden of 14.35 mutations per year per cell (Figure SI 7B), and used as rate *µ*_*b*_ = 14.35 − *µ*_*d*_ /*τ*.

**Table 1.**
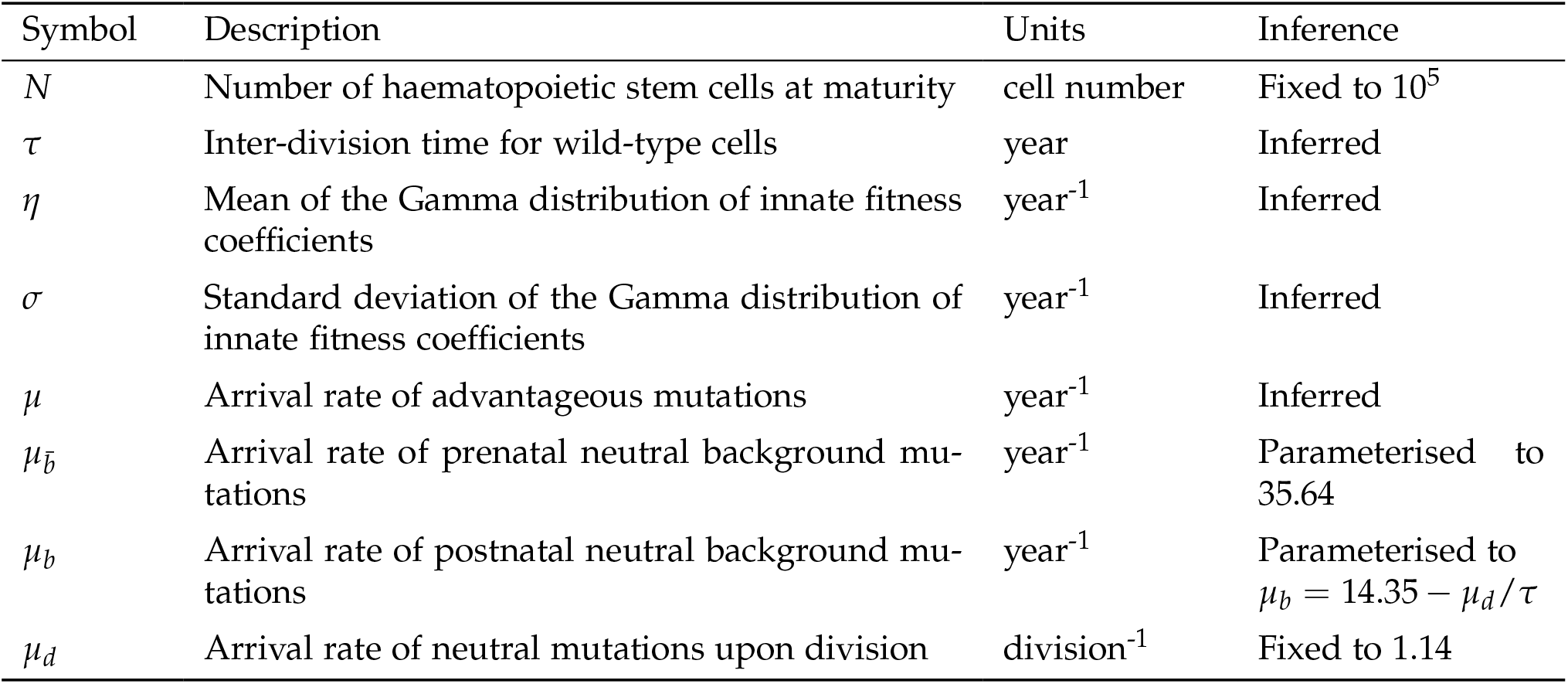
Parameters of the agent-based model. Parameters were inferred either through the ABC framework described in SI 5.4 or by regression of the mean of the number of mutations as described in the text.

Estimating the mutation rates in this manner ensures that our model accurately captures the mean of the burden in data. However, we find that it underestimates the variance, as similarly reported in previous work [12], with an error that increases with age. This suggests a high heterogeneity across cells in the number of mutations due to other processes not modelled in our framework (Figure SI 7C-D).

### SI 5.4 Bayesian inference on site frequency spectra to estimate the fitness distribution

We then investigated the identifiability of the underlying evolutionary parameters of CH from *in vivo* single-cell mutational data. We first considered the qualitative behaviour of the clonal competition model based on values of the arrival rate of advantageous clones *µ*, and the mean *η* and standard deviation *σ* of the innate fitness distribution. Faster dynamics can be obtained by higher values of *η* and *µ*, and lower values of *τ* (Figure SI 8).

To test what combinations of these parameters { *τ, µ, η, σ* } best describe the data, we implemented approximate Bayesian computation (ABC) inference schemes [20]. ABC is a Monte Carlo likelihood-free method that samples different parameter sets and stores those generating the simulations which are close to the data according to a metric of similarity.

#### SI 5.4.1 Summary statistics

We implemented ABC by computing two summary statistics: the SFS *S* and the number of clones *C* with frequency greater than 1%. We quantified the similarity between the data *y* and the simulations *z* by computing the Wasserstein distance for the SFS *w*(*S*_*y*_, *S*_*z*_) and the relative Euclidean distance for the number of detected clones *d*(*C*_*z*_, *C*_*y*_) = | *C*_*z*_ − *C*_*y*_ | /*C*_*z*_. The Wasserstein distance, also called the Earth mover’s distance or the optimal transport distance, is a similarity metric between two probability distributions which computes the unsigned average distance between the inverse cumulative distribution of *S*_*z*_ and *S*_*y*_ [21].

#### SI 5.4.2 Longitudinal inference using the SFS and the number of clones

We performed a longitudinal inference to estimate the values of { *τ, µ, η, σ*} from the single-colony data published in Mitchell et al. [4]. As mentioned above, the data consist in *in vitro* expanded single-cell colonies collected from either the bone marrow, the peripheral or cord blood of 9 healthy donors aged 0 to 81. Since the individuals are healthy donors, we assumed that they followed the same genetic process, i.e. we assumed that the individuals represented timepoints of the same underlying stochastic process. Being interested in the evolution of the CH dynamics over time, we needed exactly one datapoint per age: we kept the newborn with the higher number of cells and discarded the other one, running the inferential task on 8 donors in total.

We generated 65,000 realisations of our agent-based model on the high-performance cluster [22] drawing different sets of parameters from uniform prior distributions. We saved the state of the simulations after subsampling the stem cell population to the number of cells found in the 8 donors at their respective ages (i.e. age 0, 29, 38, 48, 63, 76, 77, 81). We then quantified the similarity between the simulations *z* and data *y* using the two summary statistics described above. We obtained the posterior distributions by storing the parameter sets that generated simulations with both metric distances within the 1% quantile for at least 7 out of 8 timepoints *i*, i.e. the parameter sets for which the simulations had 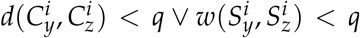, with *q* being the 1% quantile. These filtered parameters are approximated samples of the posterior distributions that describe the data best. We summarised the posterior distributions into point estimates by taking their modes after binning them into histograms, reporting the 90% credible interval. The summary statistics for the 8 donors are shown in Figure SI 9 and Figure 3D. The simulations that minimise 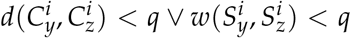 recapitulate the stochastic behaviour of clonal expansions, detected in the SFS as peaks in the higher frequency domain (Figure SI 10).

The inferred posterior distributions show estimates of mean and standard deviation per year of 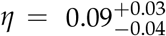 and 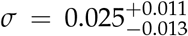, and an arrival rate of advantageous variants of 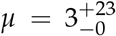 (Figure SI 11). Taking the point estimates of the posterior distributions of *η* and *σ* allows us to plot the Gamma distribution of the fitness effects inferred from the data by mapping *η* and *σ* to the scale *α* and shape *θ*: *α* = *η*^2^/*σ*^2^ and *θ* = *σ*^2^/*η*. The recovered distribution of fitness effects is bell-shaped, suggesting that most clones expand at similar rates (Figure 3K). Moreover, we find that fit clones expand slowly, with an average growth that is 9% faster per year compared to wild-type cells. The findings do not depend on the number of runs accepted by the Bayesian framework, the estimates being consistent across a range of accepted runs of 0.1%, 0.5% and 5% (Figure SI 12).

#### SI 5.4.3 Testing the Bayesian framework on simulated data

To validate our inferential framework, we tested our implementation on synthetic data generated by the ABM simulations with *η* = 0.10, *σ* = 0.723, *µ* = 4.86 and *τ* = 1.16. We found 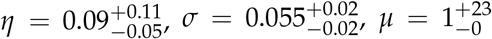 and 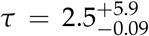 (Figure SI 13), confirming the validity of the inferential framework for this parameter range.

## SI 6 Combining three longitudinal sequencing datasets

To apply the clonal competition model to a large dataset, we aggregated the target sequencing data generated by Fabre et al. [2], by Robertson et al. [10] and by Van Zeventer et al. [23]. These are three longitudinal studies of DNA blood samples, collected from a total of 1,969 individuals with no history of haematological malignancy, for which a target panel of driver genes was sequenced. Some donors in the Van Zeventer et al. [23] had anaemia, cytopenia and others peripheral blood count abnormalities. Variants with a frequency smaller than 1% were removed from Fabre et al. and from Van Zeventer et al. A stricter threshold of frequency smaller than 4% was adopted for Robertson et al., resulting in 1,923 viable donors across all data sets.

Clonal haematopoiesis of indeterminate potential (CHIP) is defined by the presence of at least one expanded clone with VAF ≥ 2% [24]. To estimate the prevalence of CHIP in the data, we stored the donors’ age at which the first variant with a frequency greater than 4% occurred. These ages were binned into two-year spans, generating a distribution of ages. Then we took the cumulative sum of this distribution, and normalised it by the total number of donors (with and without CHIP) to obtain the prevalence. We found 727 out of 1,923 total donors (37.81%) without any evidence of CHIP (Figure SI 15B, light blue), possibly underestimated by some missed clones. Half of the donors had CHIP by 75 years of age, showing that CHIP is prominent in healthy individuals. The prevalence of CHIP correlates with age (dark blue), as expected, and shows three different regimes: initially, it increases linearly, then sublinearly, and finally stays constant in the elderly. As most of the datapoints lie between 60-92 years (Figure SI 15A), the initial linear phase and the constant final phase might be biased by a small sample size.

To mitigate the age sampling bias in estimating the prevalence if CHIP, we computed the rate of CHIP incidence. Unlike the prevalence of CHIP, which is calculated by dividing the cumulative age distribution by the total number of donors, here we normalised the number of donors with CHIP within two-year age bins by the total number of donors in each bin. The estimated CHIP rate peaks around 60 years and declines at older ages (Figure SI 15C). This observed peak is likely influenced by age bias in the dataset, as very few data points were collected from individuals younger than 60 years. If blood samples from younger donors had been available, a more gradual increase between 55 and 60 years would likely have been observed. Additionally, it is possible that targeted sequencing approaches missed smaller clones that had already expanded at an earlier stage. Finally, the decrease in CHIP rate can be explained by the survival bias present in this dataset of healthy individuals, as cancer patients, who usually show large clones at younger ages, were excluded.

Plotting the prevalence of some frequently mutated genes showed that mutations were common in epigenetic regulator genes *DNMT3A* and *TET2*, and also frequent in *ASXL1, TP53, PPM1D* (Figure SI 15D). Figures SI 15E-G show the changes in VAF over time for *JAK2*-V617F, *DNMT3A*-R882H and *DNMT3A*-R882H, where each line corresponds to a donor (trajectory). *JAK2*-V617F was the most prevalent variant found in 67 donors, followed by *DNMT3A*-R882H, found in 49 donors, and *SRSF2*-P95H, found in 26 donors. Sequencing only 79 genes already shows a complex fitness landscape according to which trajectories can stay constant, increase or decrease over time. This suggests a dynamic fitness landscape that can be recapitulated by the clonal competition model, as opposed to simpler models that fail to do so. Interestingly, the data show that the same variant can have different growth effects in donors, suggesting that the fitness advantage conferred by the variant depends on the context in which it occurs, influenced by the presence of other clones, the microenvironment or other epigenetic factors.

## SI 7 Bayesian inference of innate fitness and arrival time on individual trajectories

To estimate the innate fitness 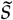 and arrival time 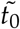 of an observed target trajectory we performed an approximate Bayesian computation comparing it to simulated reference trajectories using the SDE model (1) with the best-fit parameters obtained from the ABC described in SI 3. Each reference trajectory was generated by inserting one single-celled clone with a predetermined fitness *s*_*i*_ at an arrival time *t*_*i*_ in a single realization of the SDE and tracking it over time. Drawing all *s*_*i*_ and *t*_*i*_ from uniform prior distributions *s* ∼ *U*(0.01, 2) and *t* ∼ *U*(0, 90) we obtained an ensemble of 150,000 independently simulated trajectories. For a target trajectory 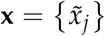 measured at ages { *z*_*j*_ }, the distance from a reference trajectory *x*_*i*_(*t*) was c{alculat}ed as the Euclidean distance 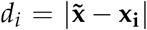 from the reference measured at the same times **x**_**i**_ = *x*_*i*_(*z*_*j*_) . To generate the posterior distribution we retained only trajectories with distances up to the 150th order statistic (0.1% of all trajectories). Finally, to obtain estimators for 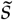 and 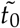 we performed a kernel density estimate of the marginalized posteriors and took their maxima (main text Figure 4B).

### SI 7.1 Estimation error arises from inherent stochasticity

To probe the accuracy of this method we generated ensembles of target trajectories with a fixed parameter, for example a fixed 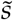 and random 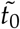, or vice versa. We then performed the ABC comparison on each trajectory in the ensemble, resulting in a distribution of estimates for the fixed parameter (main text Figure 4C). This distribution encodes the inherent uncertainty of the inference method, which arises from the randomness of the evolutionary process itself. Indeed, without specific knowledge of competing variant behaviours, a variant originating at *t*_0_ with fitness *s* has a distribution of possible future trajectories, all of which have a certain likelihood of occurring. By running many realizations of the process, the ABC reproduces this distribution for all possible (*t*_0_, *s*) combinations. Upon comparison with a target trajectory, it identifies the (*t*_0_, *s*) pair with the most similar expected trajectory. Thus variants which evolve according to high likelihood scenarios (i.e. close to their expected trajectory) will have trajectories similar to mean trajectories from nearby points in parameter space. However a variant which ends up on an “unlikely” trajectory, far from its expected value, will be identified as closer to a more distant parameter pair 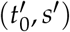. Overall, we found the errors to be unbiased for wide ranges of *s* and *t*_0_, excepting very early arrival times (*t*_0_ ≈ 0), which had a slight positive bias. The error distributions also showed slight skewing depending on the parameter value, and increasing variance for higher fitnesses and lower arrival times.

### SI 7.2 Genic variant fitness estimates from multiple donors

To estimate the innate fitness of a particular variant for which we have multiple trajectories (e.g. occurrence in multiple donors), we note that a single trajectory fitness estimate *ŝ* is subject to noise arising from the above-described error, as well any other factors, such as measurement errors or potentially unconsidered processes. We thus write

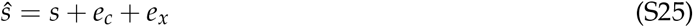

where *s* is the variant’s true innate fitness and we have split the noise into a contribution from the Bayesian estimation *e*_*c*_ due to stochastic competition (see SI 7.1) and a contribution from all other sources *e*_*x*_. Since *e*_*c*_ is unbiased, and assuming *e*_*x*_ is as well, we have ⟨*ŝ*⟩ = ⟨*s* + *e*_*c*_ + *e*_*x*_⟩ = *s* and we can take the mean of the trajectories as estimator for the true innate fitness. It has a standard error that depends on the number of trajectories *n* as 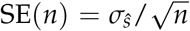 is the standard deviation of the distribution of individual estimates. This can be found from the variance, which – further assuming the errors are not correlated – takes the form

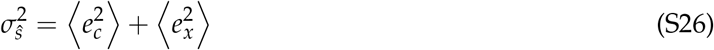

Here 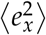 is by definition unknown, as it describes all error contributions that were not modelled. We can however obtain *σ*_*c*_ by performing the Bayesian inference on a large set of simulated trajectories (with identical innate fitness). In this case *e*_*x*_ = 0 and taking the variance of the fitness estimates directly gives 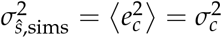. For an innate fitness of *s* = 0.15 this resulted in *σ*_*c*_ = 0.026 (main text Figure 4G). To estimate the total variance we then looked to the data. Considering the sets of fitness estimates per variant {*s*_*i*_ }_var_ as independent samples, we can interpret their respective standard deviations 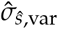 as independent estimators of *σ*_*ŝ*_. Of the variants considered (> 10 occurrences in the donor data), DNMT3A p.R882H/C and JAK2 p.V617F were the highest recurring, with n=47 and n=46 respectively (compared to n=16 for the next highest), and thus their standard deviations of 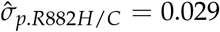 and 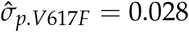 have the lowest statistical error with regards to the true *σ*_*ŝ*_. Furthermore, because each variant represents an independent sample population, the average over all standard deviations (one from each variant) acts as a bootstrapped estimator for *σ*_*ŝ*_, for which we find 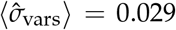. Given this agreement we take *σ*_*ŝ*_ = 0.029, and thus for each variant

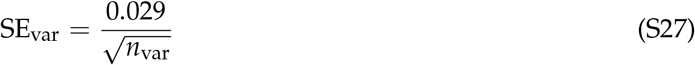

For the most frequently occurring variants (*n* > 10) we obtained the fitness estimates shown in Table 2. Given estimates of *σ*_*ŝ*_ and *σ*_*c*_, we can conclude that competition explains 80% of the total variance 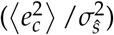 observed in trajectories of identical variants.

## SI 8 The arrival time distribution of detectable variants

In order to evaluate the distribution of arrival times estimated from donor variant trajectories we investigate model predictions for this quantity.

### SI 8.1 Single event clones

We first consider the case of variants arriving randomly in time according to the Poisson process, consistent with the observation of linearly accumulating mutations in the population. Denote *S*_*T*_(*t*) the expected number of detectable variants measured at time *T* which originated at time *t*. To obtain an expression for this we consider the following simplifications of the previously described model (SI 2):

- Deterministic dynamics: newly arising clones grow according to a fixed growth function, i.e. experience no fluctuations and do not go extinct.
- Exponentially growing clones: we take clones to grow independent of any compartmental or competitive limitations, as we are only interested in the early growth period, until a clone reaches the detection threshold.

**Table 2.** Inferred innate fitness of commonly occurring variants.

| Variant | Trajectories | $s_{innate}$ | 2SE |
| --- | --- | --- | --- |
| DNMT3A_p.P904L | 13 | 0.14 | 0.016 |
| DNMT3A_p.Y735C | 13 | 0.14 | 0.016 |
| DNMT3A_p.R882H/C | 47 | 0.15 | 0.0085 |
| TET2_p.I1873T | 14 | 0.15 | 0.016 |
| JAK2_p.V617F | 46 | 0.16 | 0.0086 |
| ASXL1_p.G646fs | 13 | 0.16 | 0.016 |
| SF3B1_p.K700E | 11 | 0.14 | 0.017 |
| SF3B1_p.K666N | 12 | 0.17 | 0.017 |
| SRSF2_p.P95H13 | 16 | 0.18 | 0.015 |

Under these assumptions, with *µ* the arrival rate of new clones and *η* and *σ*^2^ the mean and variance of the (Gamma-distributed) fitness distribution, the expected number of detectable variants is given by:

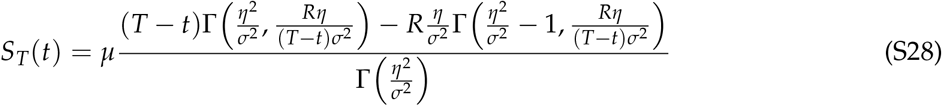

with *R* = log(*Z*/*n*_0_) where *Z* is the size threshold for detection and *n*_0_ is the starting size of newly initiated clone.

To derive this we first write *S*_*T*_(*t*) as an integration over time and fitness:

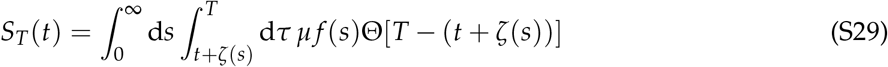

where Θ(*x*) is the Heaviside step function, and *ζ*(*s*) is the time required for a clone to grow from its inception size *n*_0_ to the detection threshold *Z*. This expression essentially states that the total number of detectable clones can be found by integrating over contributions from clones at all fitnesses. The contribution from clones at a particular fitness is found by integrating the constant arrival rate over the timespan from when they would first become detectable (given their arrival at time *t*) until the measurement time. The step function appears because we must only count clones that can actually reach the detection threshold within the given time frame *T* − *t*. Under deterministic non-competitive dynamics clones grow exponentially, so that *ζ*(*s*) = log (*Z*/*n*_0_)/*s* = *R*/*s*. The inner integral can be performed immediately. The step function can be absorbed into the lower bound of the second integral (over *s*) with the assumption that *ζ*(*s*) < *T* − *t*. Plugging in the probability density function of the Gamma distribution with shape *α* and scale *θ* we are then left with

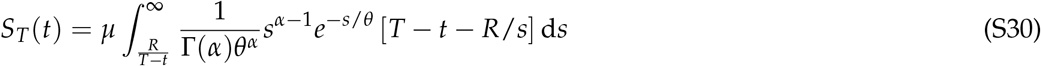

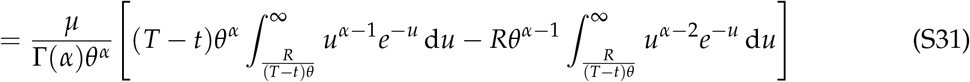

Where in the second equality we have performed the integral substitution *u* = *s*/*θ*. Here we recognize the incomplete gamma function

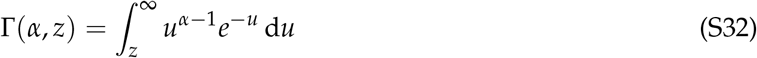

in the first term and thus Γ(*α* − 1, *R*/(*T* − *t*)*θ*) in the second. Finally, substituting shape and scale for mean and variance through the relations *α* = *η*^2^/*σ*^2^ and *θ* = *σ*^2^/*η* we obtain the result in Equation (S28).

### SI 8.2 Multiple-event clones

To obtain a prediction for the arrival time distribution of clones originating with epistatic interactions, e.g. a second or third mutation, we modified our simulations (see SI 2.1) to include compounding fitness effects in the following manner. For each newly arising fit variant, a parent clone is chosen randomly from all currently existing clones in the system. The likelihood of a parent being selected is given by its population fraction. If the parent is the wild type, then the clone receives an innate fitness randomly drawn from the fitness distribution, just as in the non-epistatic model. If the parent is an expanding driver, the new clone receives a fitness that is a combination of the parent’s innate fitness and a value drawn from the fitness distribution. We used a multiplicative fitness to model the epistatic effect of multiple variants:

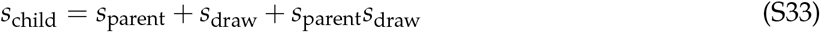

where *s*_child_ is the innate fitness of the newly arising clone, *s*_parent_ the innate fitness of the parent, and *s*_draw_ the fitness granted by the novel mutation.

We then ran a large number of simulations and recorded the arrival times of all detectable clones (threshold 0.01%) with at least one driver measured at random times between ages 60 and 95, resulting in the age distribution shown in Figure 4.

---

**Figure SI 1.**
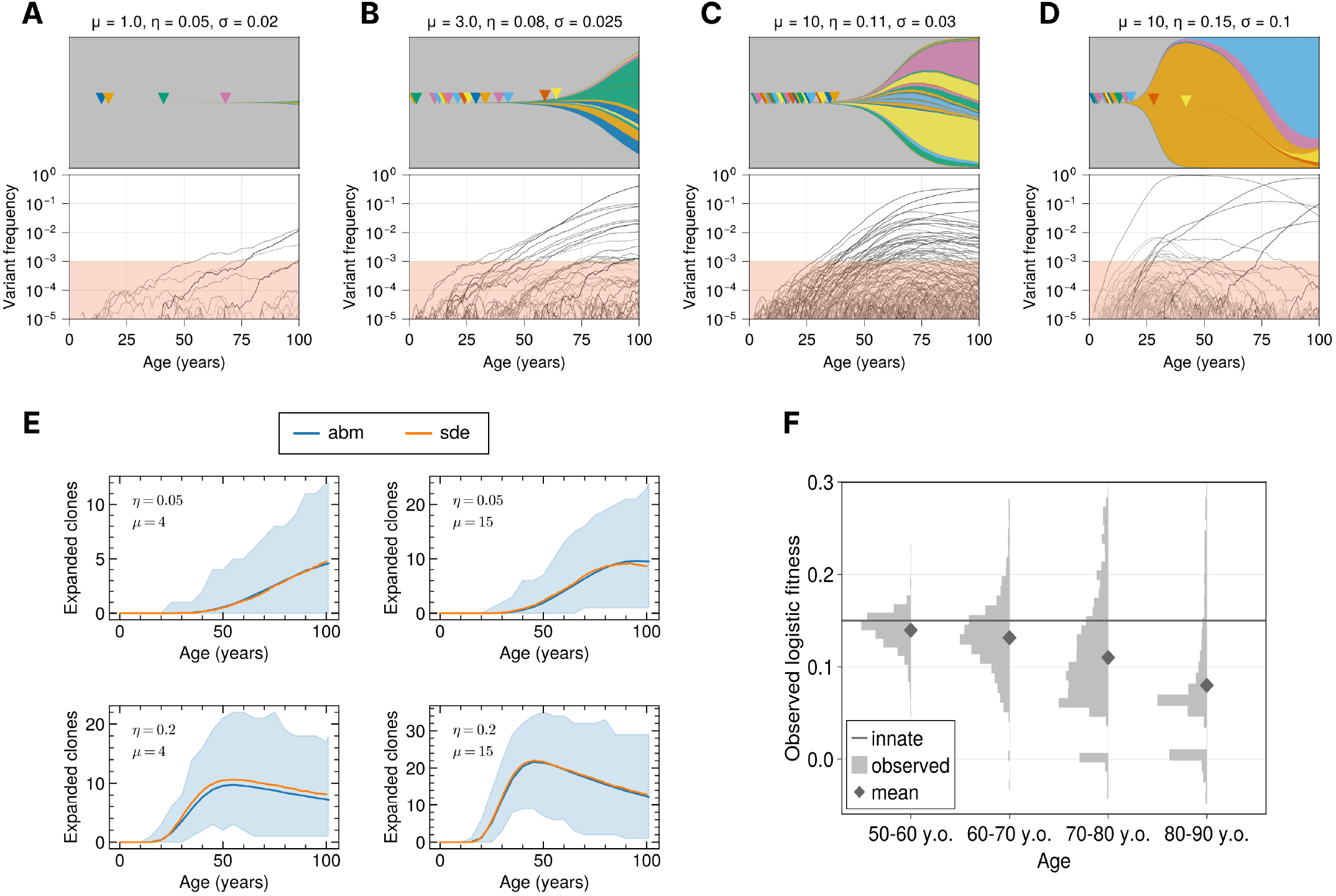
Model dynamics testing and exploration. **A-D)** Example realisations of the SDE model (Equation (S19)) for different parameter combinations. In the Muller plots (top row) the mutant arrival times are indicated by triangles. **E)** The number of detectably expanded (>0.5%) clones with age averaged over 500 simulations agrees between the SDE (SI 2.1) and ABM (SI 4.2) implementations of the model. The blue shaded area shows the minimum and maximal values of the expanded clones for the ABM model. **F)** In 10’000 simulations a variant with innate fitness *s* = 0.15 is introduced at age 10 and observed at different time points later in life. The distribution of measured logistic fitnesses becomes wider and its mean decreases with age.

**Figure SI 2.**
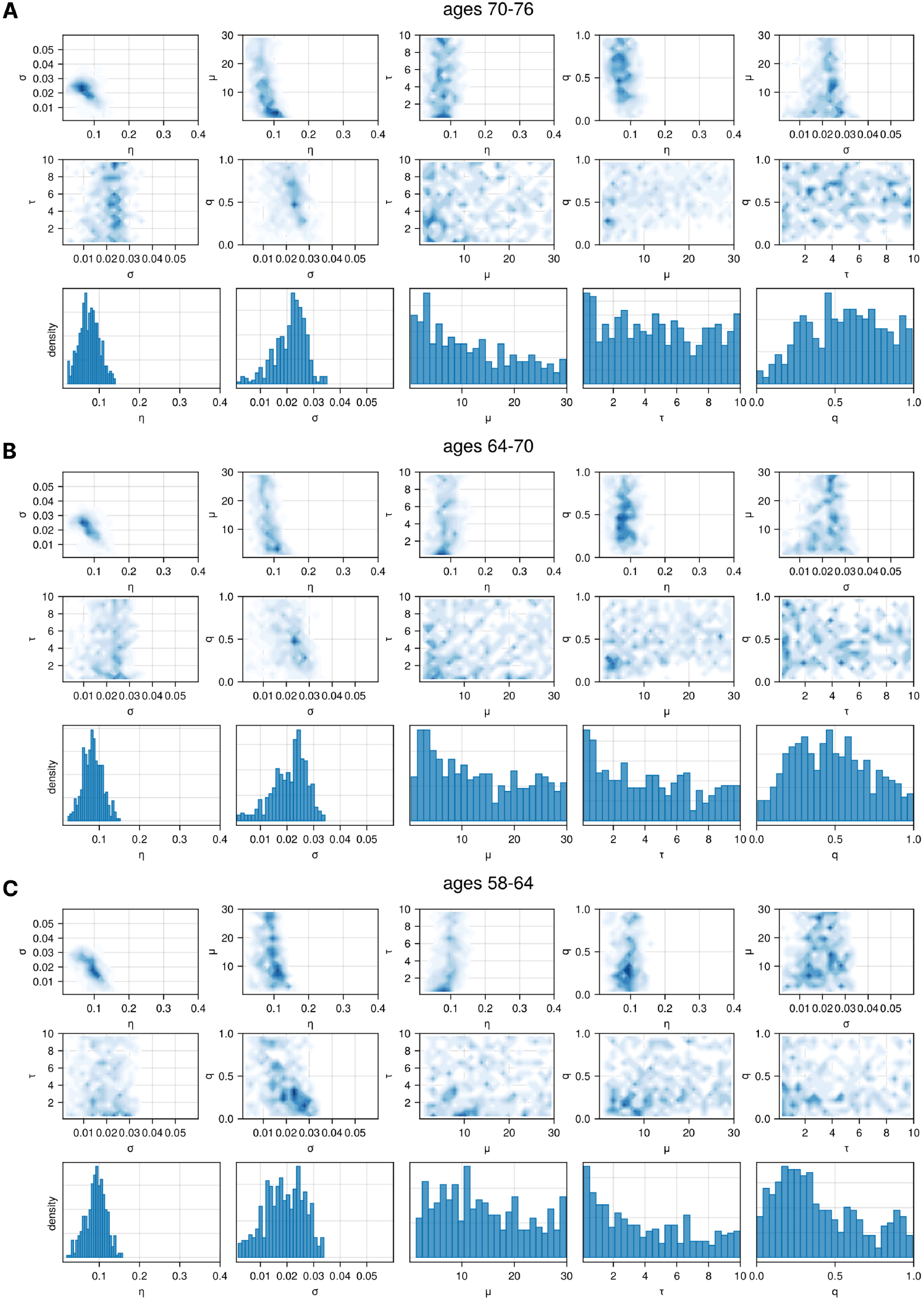
Parameter posterior distributions obtained via approximate Bayesian computation on time-resolved clone trajectories from Fabre et al. donor data. For each age group 40’000 virtual patients were simulated with parameters randomly drawn from uniform priors, with only the 1% most similar retained in the posterior. Top rows: 2D projections of the full 5-parameter posterior distribution for parameter pairs. Bottom row: marginal distributions for each parameter.

**Figure SI 3.**
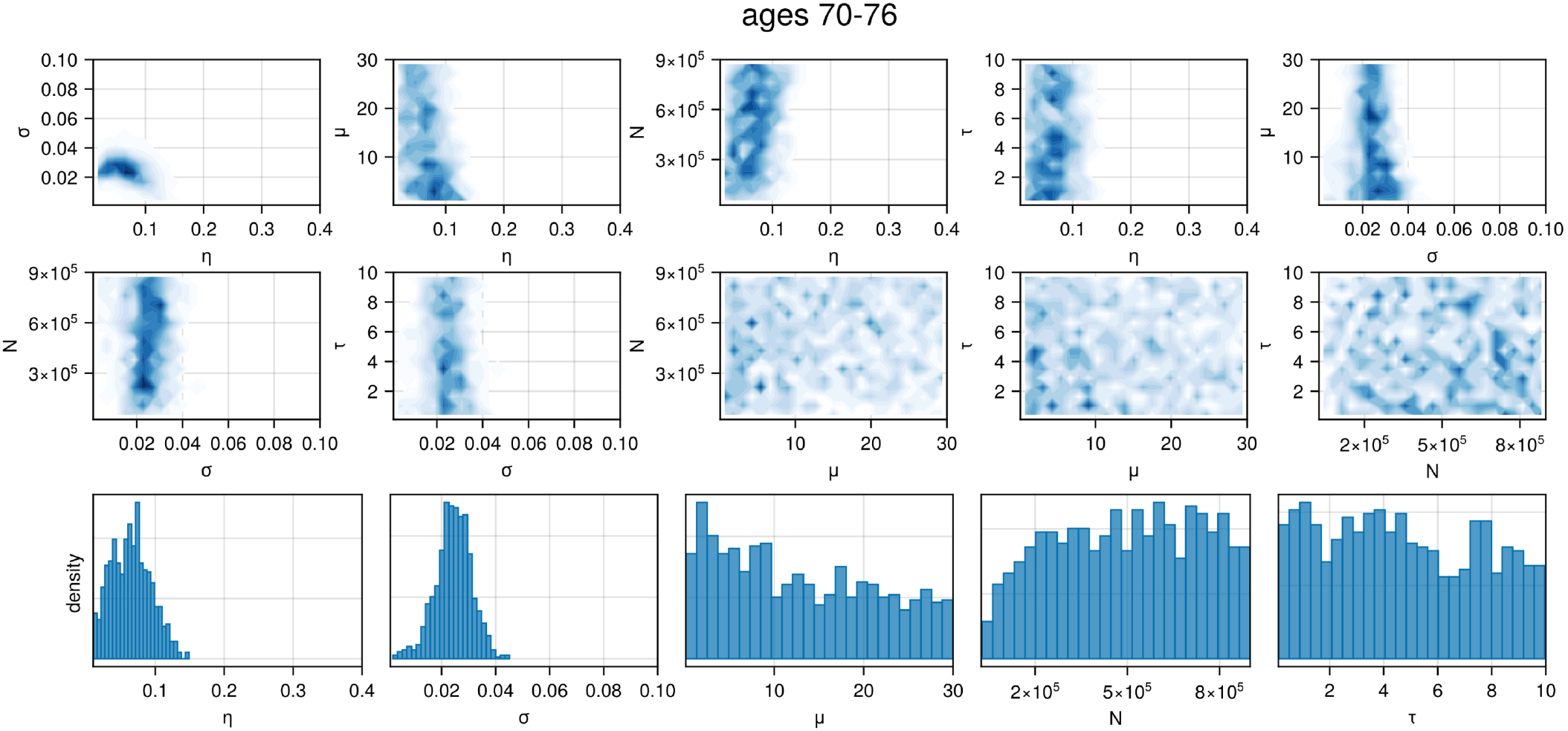
Parameter posterior distributions including the total HSC population size obtained via approximate Bayesian computation on time-resolved clone trajectories from Fabre et al. donor data. 40’000 virtual patients were simulated with parameters randomly drawn from uniform priors, with only the 1% most similar retained in the posterior. Top rows: 2D projections of the full 5-parameter posterior distribution for parameter pairs. Bottom row: marginal distributions for each parameter.

**Figure SI 4.**
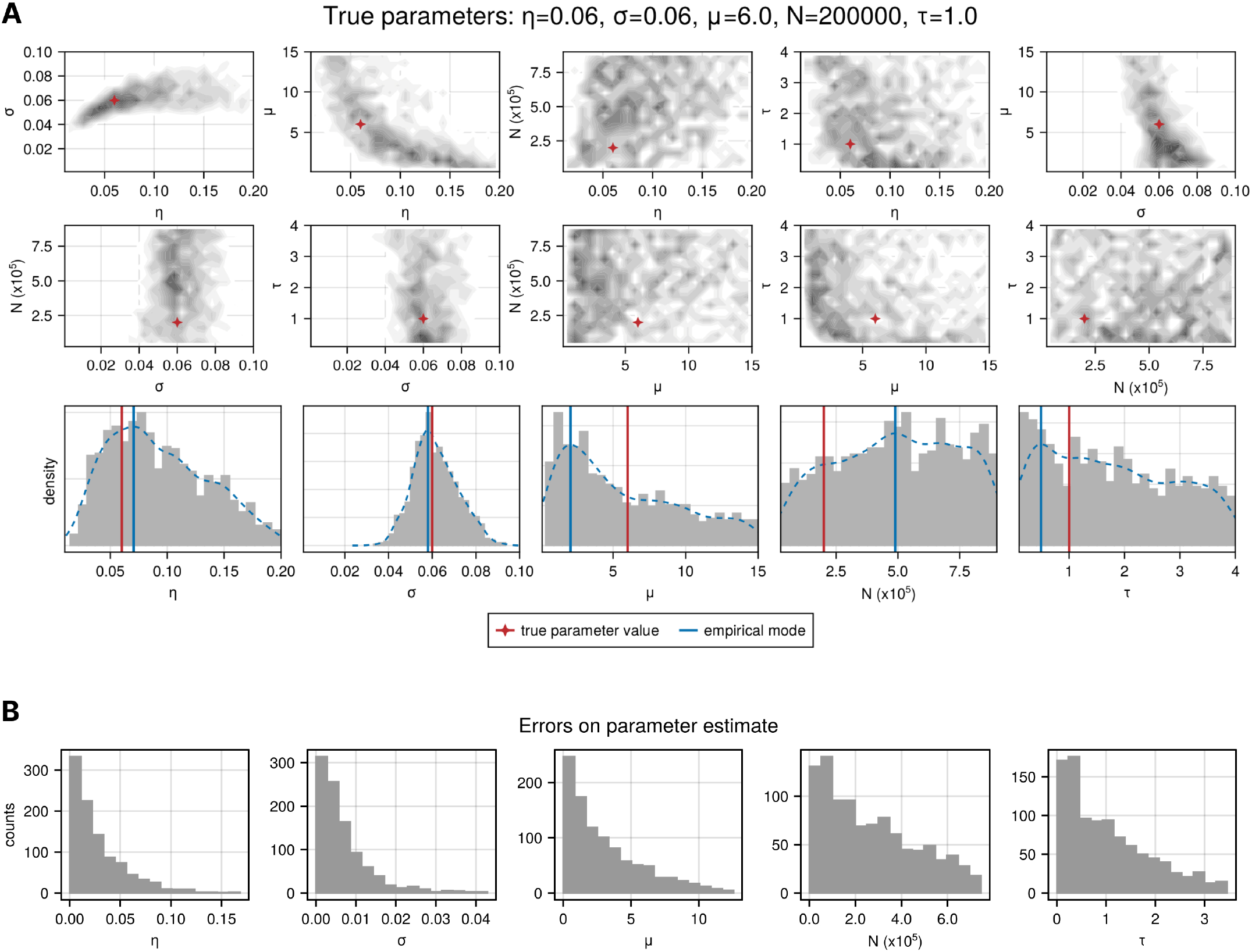
Validation of approximate Bayesian computation on variant trajectories from simulated datasets. **A)** Posterior distributions of evolutionary parameters for a single simulated dataset with known parameter values. Top rows: 2D projections of the full 5-parameter posterior distribution for parameter pairs. Bottom row: marginal distributions for each parameter, with estimated value (mode of the empirical distribution) and true value indicated as red and blue vertical lines respectively. **B)** Distribution of errors on estimated parameters from ABC for 1000 simulated datasets.

**Figure SI 5.**
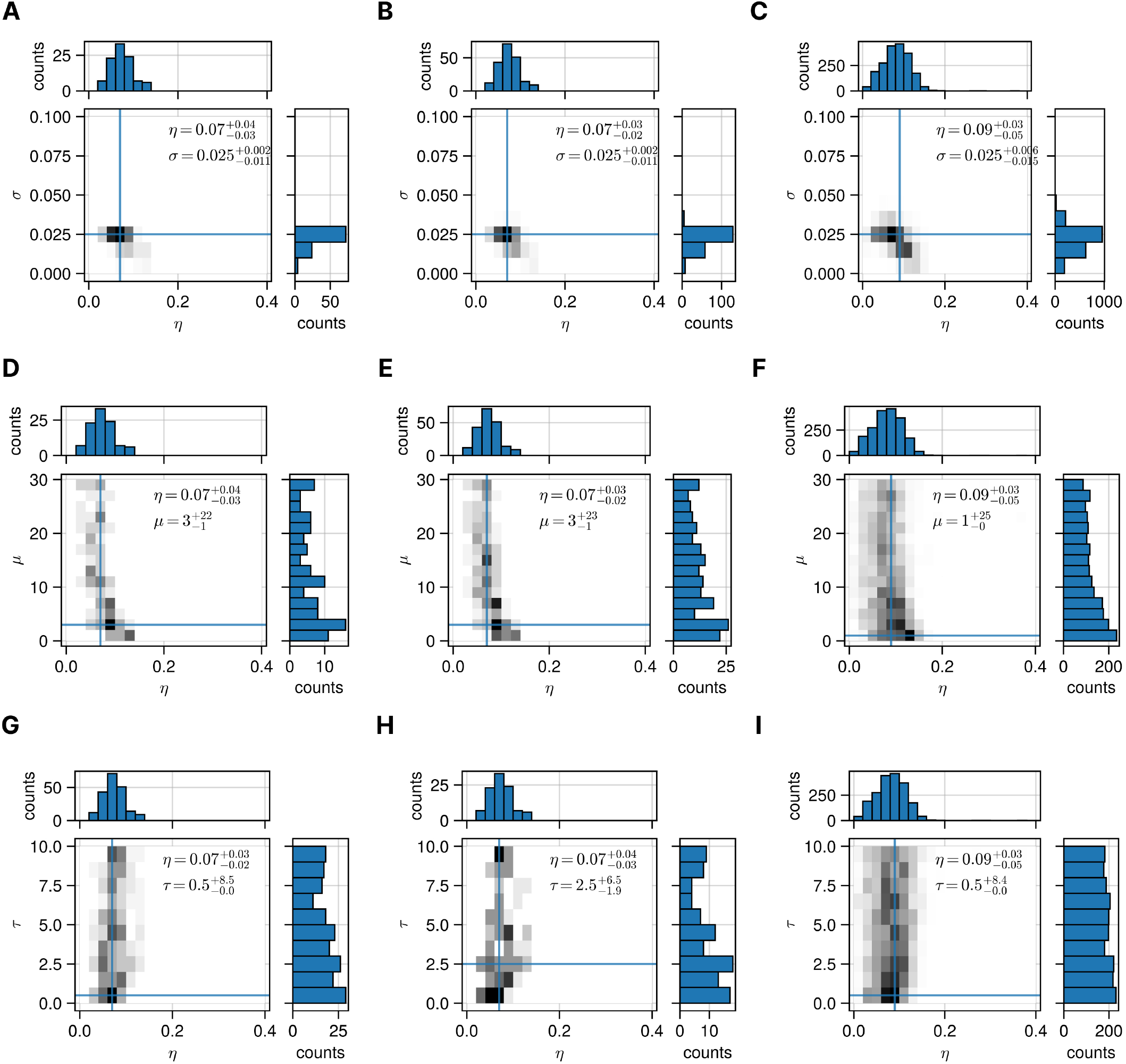
Sensitivity analysis for the posterior distributions inferred from the time-resolved clone trajectories from Fabre et al. data. The posterior distributions and their modes show consistent parameter values inferred for (*η, σ, µ*) from different percentages of the total 40’000 realisations of the HSC dynamics. This suggests that the point values inferred are not influenced by the number of samples retained. **A-C)** The 2D posterior distributions of the pair *η*-*σ* considering 0.25%, 0.5% and 5% of the runs along with the marginal posterior distributions. **D-F)** The 2D posterior distributions of the pair *η*-*µ* considering 0.25%, 0.5% and 5% of the runs along with the marginal posterior distributions. **G-I)** The 2D posterior distributions of the pair *η*-*τ* considering 0.25%, 0.5% and 5% of the runs along with the marginal posterior distributions.

**Figure SI 6.**
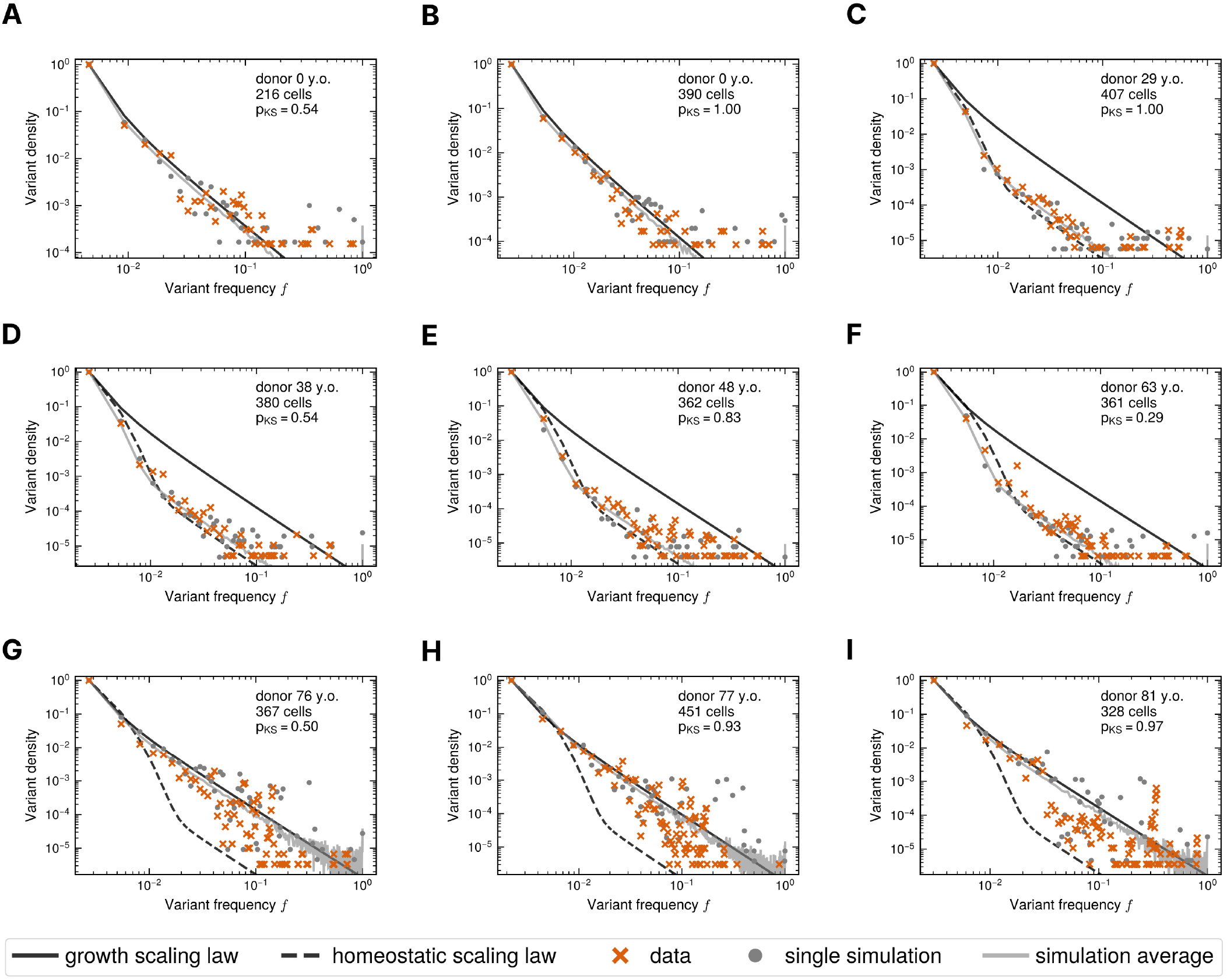
Transitions of HSC site frequency spectra (SFS) with age in Mitchell et al. data. The SFS computed from 216-451 whole genome sequenced single cells (orange crosses), from a single realisation of the agent-based simulation (grey dots), from averages over 500 simulations (grey line), from the expected scaling of a growing population (black solid line) and from the expected scaling of a homeostatic population of constant size (black dashed line). **A-B)** The SFS follow the expected scaling of a growing population for the two newborns. **C-F)** The SFS transition towards the expected scaling of a homeostatic population of constant size. **G-I)** The SFS shift back towards the expected scaling of a growing population at older ages.

**Figure SI 7.**
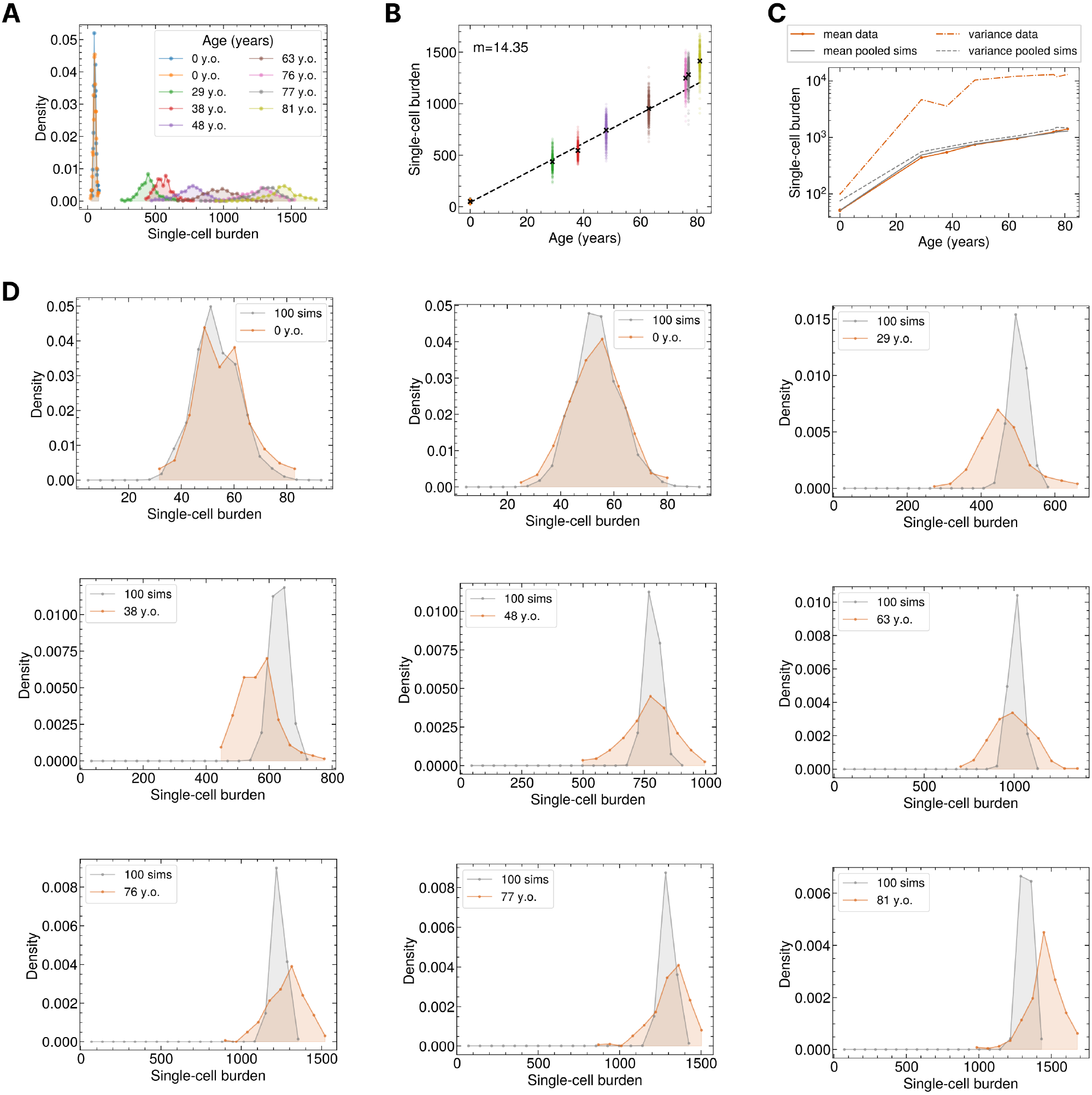
Agent-based model parametrisation with the single-cell mutational burden in Mitchell et al. data. **A)** Distribution of the number of mutations (indels and single-nucleotides) in individual cells across the donor cohort. **B)** Estimation of the average burden increase per year considering only donors without any clone at frequency greater than 1%. **C)** The mean (solid lines) and the variance (dashed lines) of the single-cell mutational burden in the data (orange) and those computed from 100 pooled simulations (grey). **D)** The single-cell mutational burden distributions in the data (orange) and those computed from 100 pooled simulations (grey).

**Figure SI 8.**
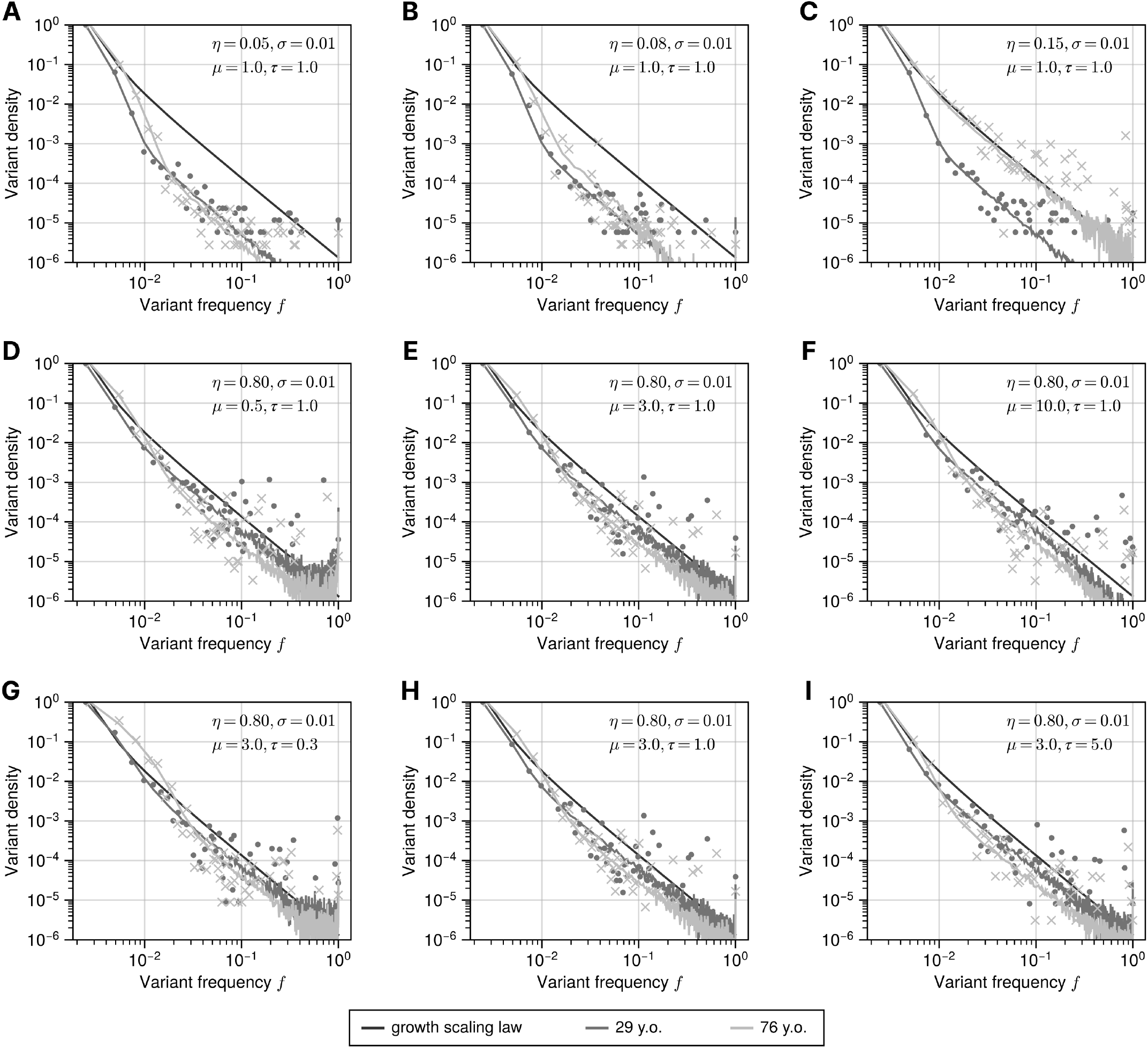
The dynamics of the simulated site frequency spectrum (SFS). Simulated SFS constructed by taking a sample of size 407 cells and 367 cells from population of 10^5^ HSCs at age 29 and 76 respectively. Grey lines show the SFS averaged over 500 simulations, dots and crosses show individual simulations, and the solid black line shows the growth scaling law considering 367 cells. **A-C)** The dynamics of the SFS varying the value of the mean of the Gamma distribution of fitness effects *η*. **D-F)** The dynamics of the SFS varying the value of the arrival rate of fit clones per year *µ*. **G-I)** The dynamics of the SFS varying the value of the inter-division time for of wild-type cells per year *τ*.

**Figure SI 9.**
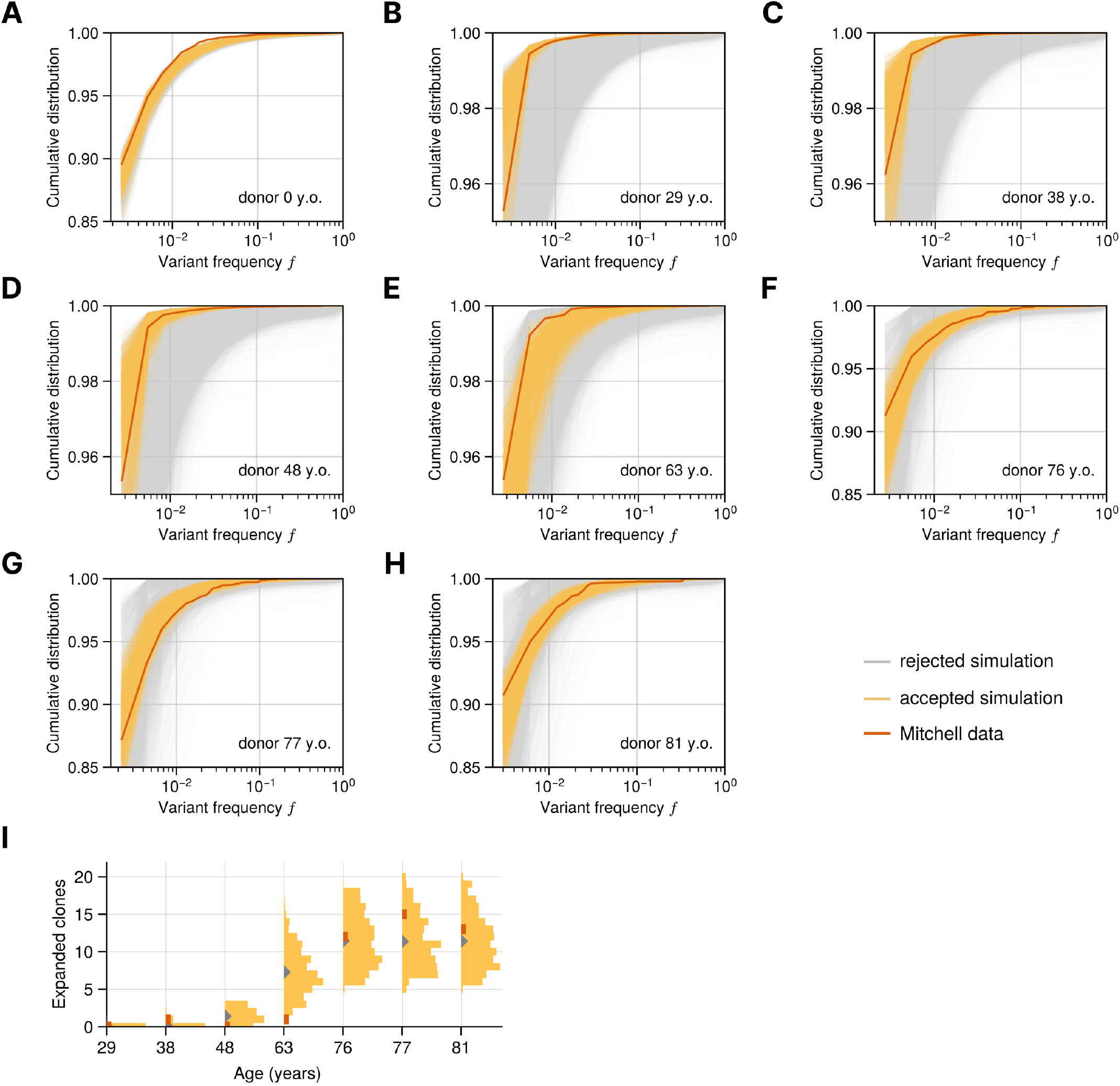
Summary statistics of the approximate Bayesian computation on the SFS and the number of detectable expanded clones from Mitchell et al. data. **A-H)** The cumulative distribution (cdf) computed from the site frequency spectra of the data (dots) is compared against the simulations. The simulations that are close to the data according to the average distance in the cdf for 7 out of 8 donors are kept (yellow) as samples of the posterior distributions. The remaining simulations (grey) are discarded. **I)** The number of clones with frequency ≥ 1% in the data (orange squares) and those in the accepted simulations (in yellow, their means are shown with grey triangles).

**Figure SI 10.**
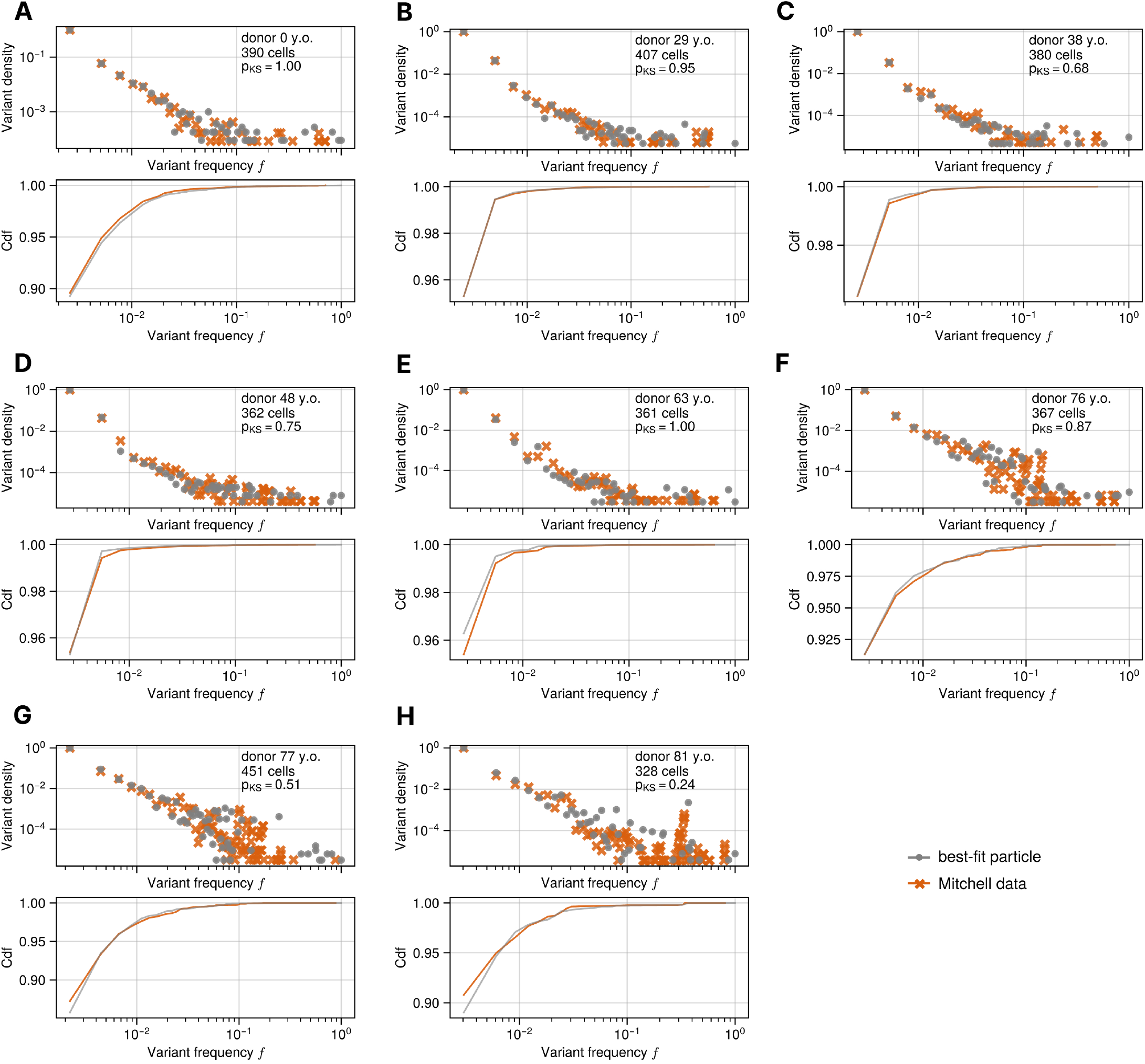
The SFS of the simulations closest to the data according to the SFS and the number of expanded clones summary statistics. The SFS (top) and the SFS cumulative distributions (bottom) computed from 216-451 whole genome sequenced single cells (orange crosses) and from the realisations of the agent-based simulation that fit the data best (grey dots). The best fits recapitulate the stochastic behaviour of clonal expansions, detected in the SFS as peaks in the higher frequency domain.

**Figure SI 11.**
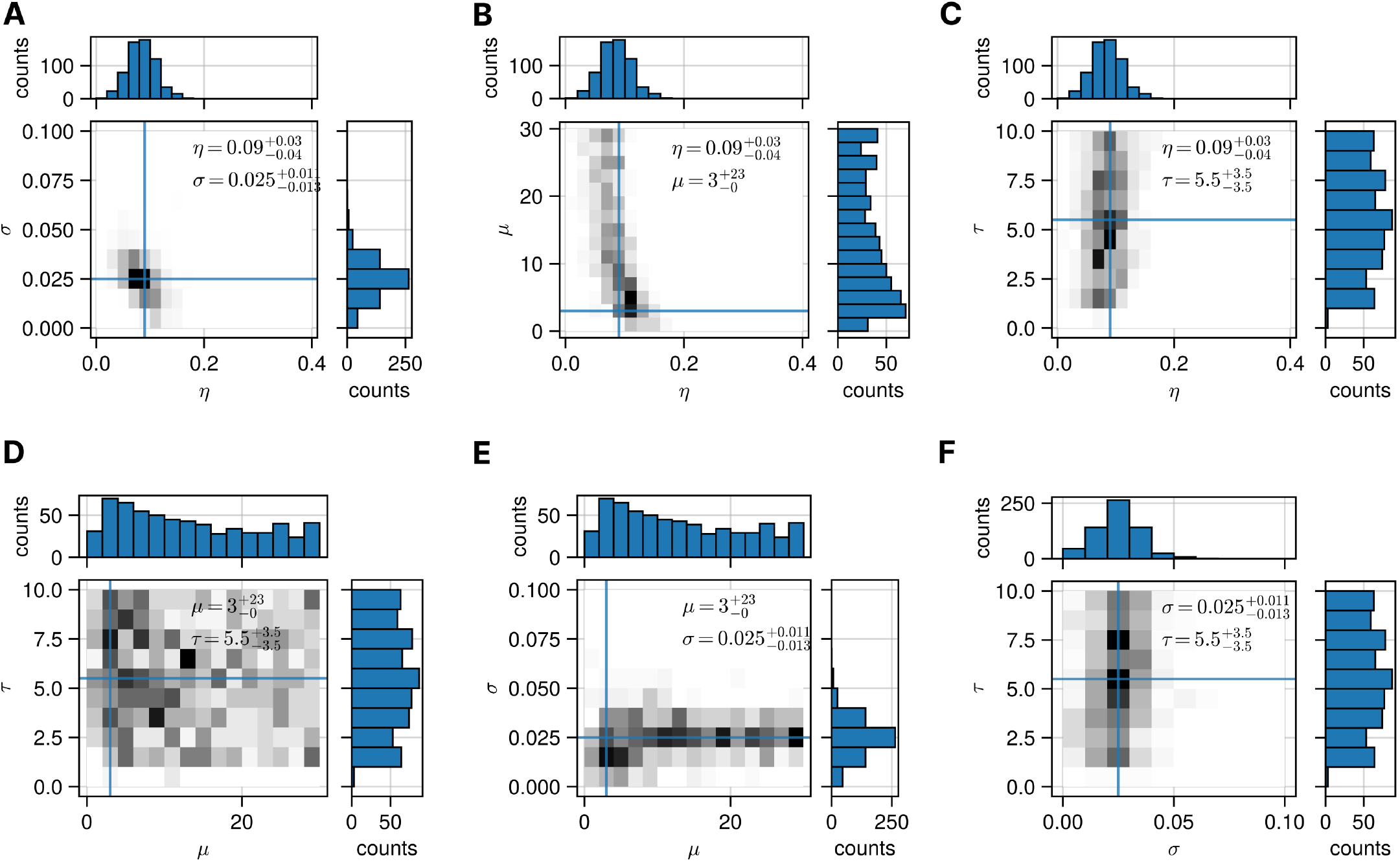
Parameter posterior distributions obtained via approximate Bayesian computation on the SFS and the number of detectable expanded clones from Mitchell et al. data. 65’000 virtual patients were simulated with parameters randomly drawn from uniform priors, with only the 1% most similar retained in the posterior. For pairs of parameters 2D projections of the posterior are shown alongside individual marginal distributions and their modes (point estimates).

**Figure SI 12.**
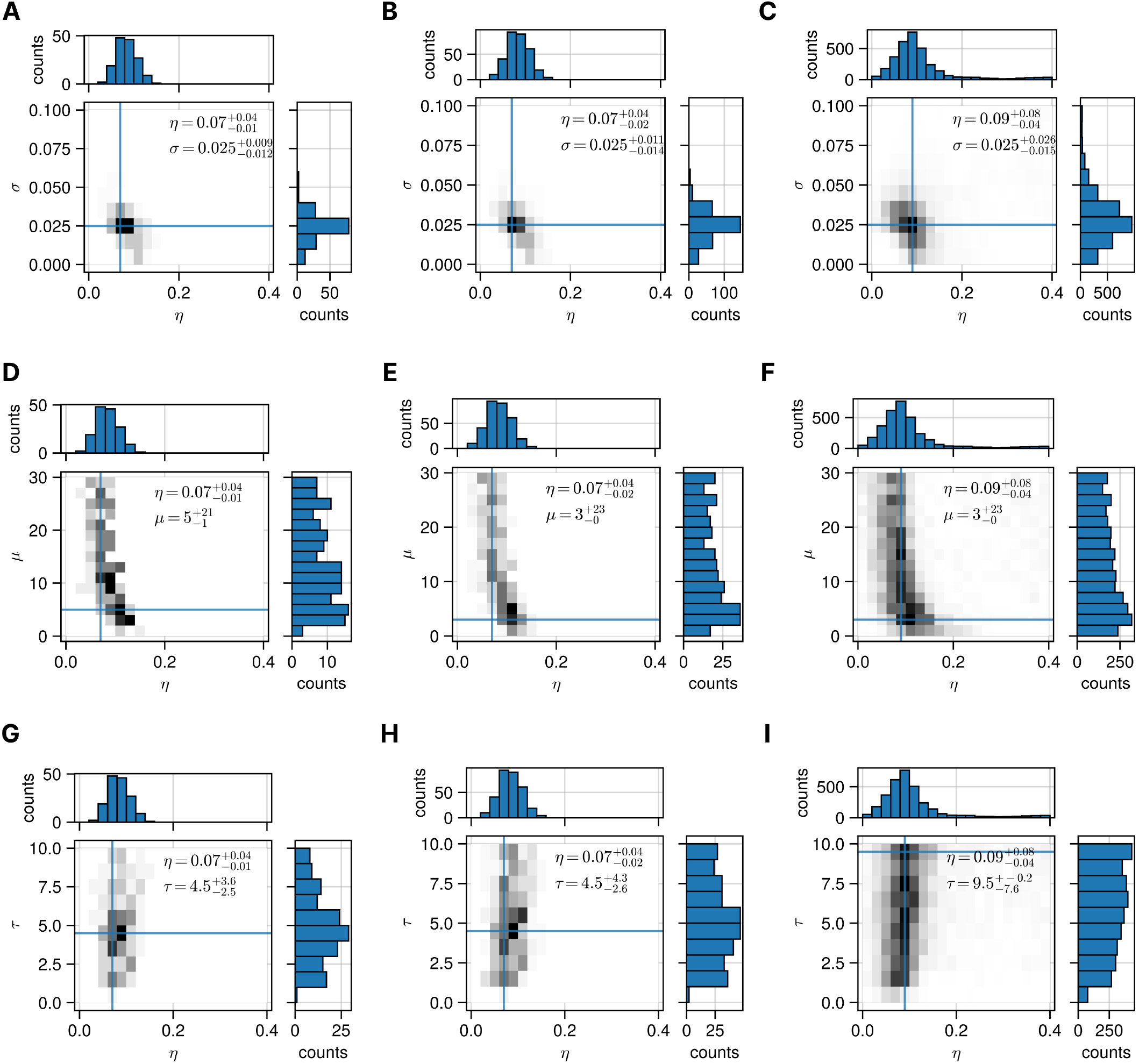
Sensitivity analysis for the posterior distributions inferred from the SFS and the number of detectable expanded clones from Mitchell et al. data. The posterior distributions and their modes show consistent parameter values inferred for (*η, σ, µ*) from different percentages of the total 65’000 realisations of the HSC dynamics. This suggests that the point values inferred are not influenced by the number of samples retained. **A-C)** The 2D posterior distributions of the pair *η*-*σ* considering 0.25%, 0.5% and 5% of the runs along with the marginal posterior distributions. **D-F)** The 2D posterior distributions of the pair *η*-*µ* considering 0.25%, 0.5% and 5% of the runs along with the marginal posterior distributions. **G-I)** The 2D posterior distributions of the pair *η*-*τ* considering 0.25%, 0.5% and 5% of the runs along with the marginal posterior distributions.

**Figure SI 13.**
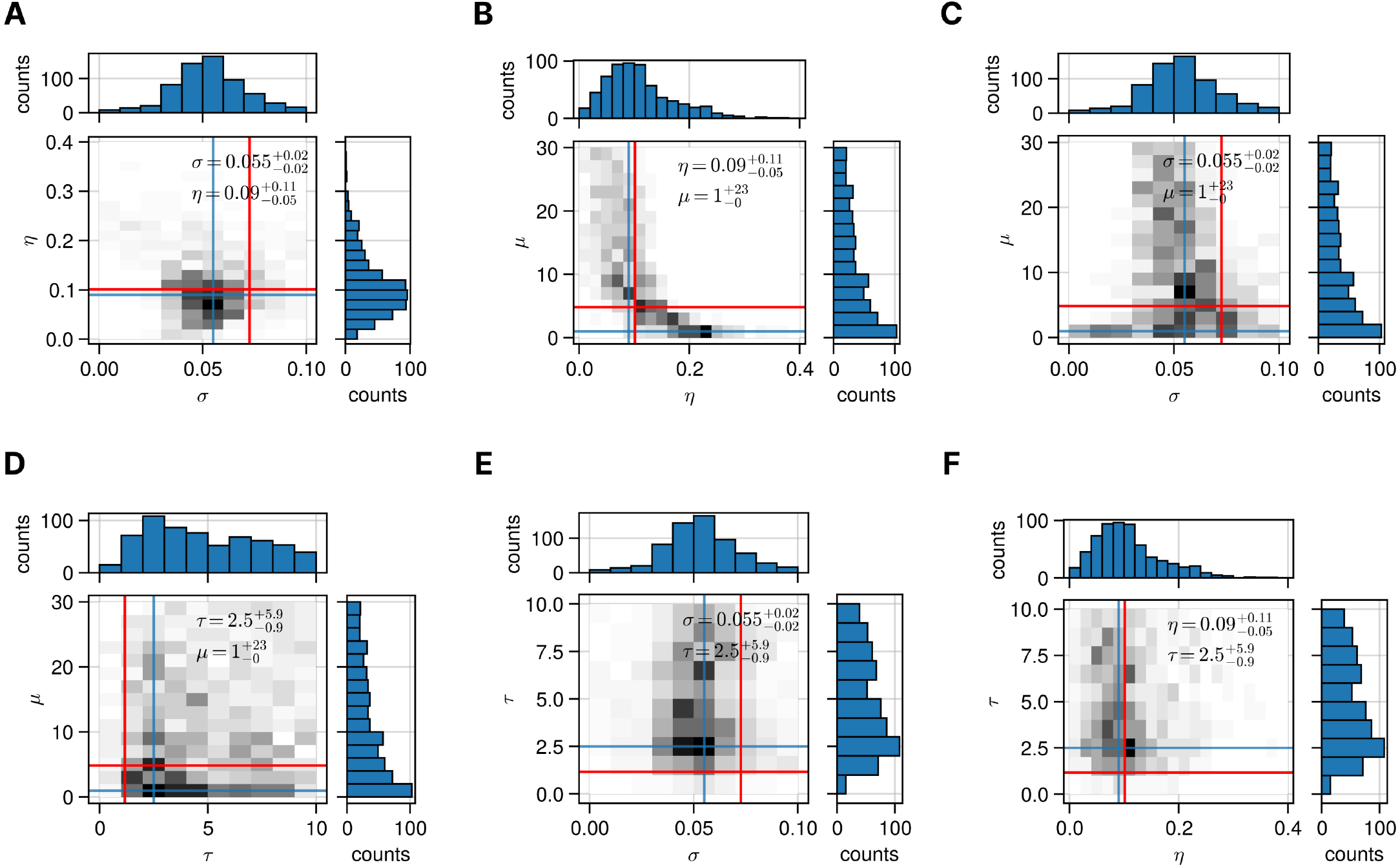
Validation of approximate Bayesian computation on SFS and number of clones on simulated data. A set of parameter values (red) have been used to generate a single realisation of the HSC dynamics, which is considered as ground truth and used to test the accuracy of the inferential framework. The posterior distributions computed from 1% of the 65’000 realisations. The inferred point estimates (modes) are shown (blue). The target values are: *η* = 0.10, *σ* = 0.072, *µ* = 4.86, *τ* = 1.16.

**Figure SI 14.**
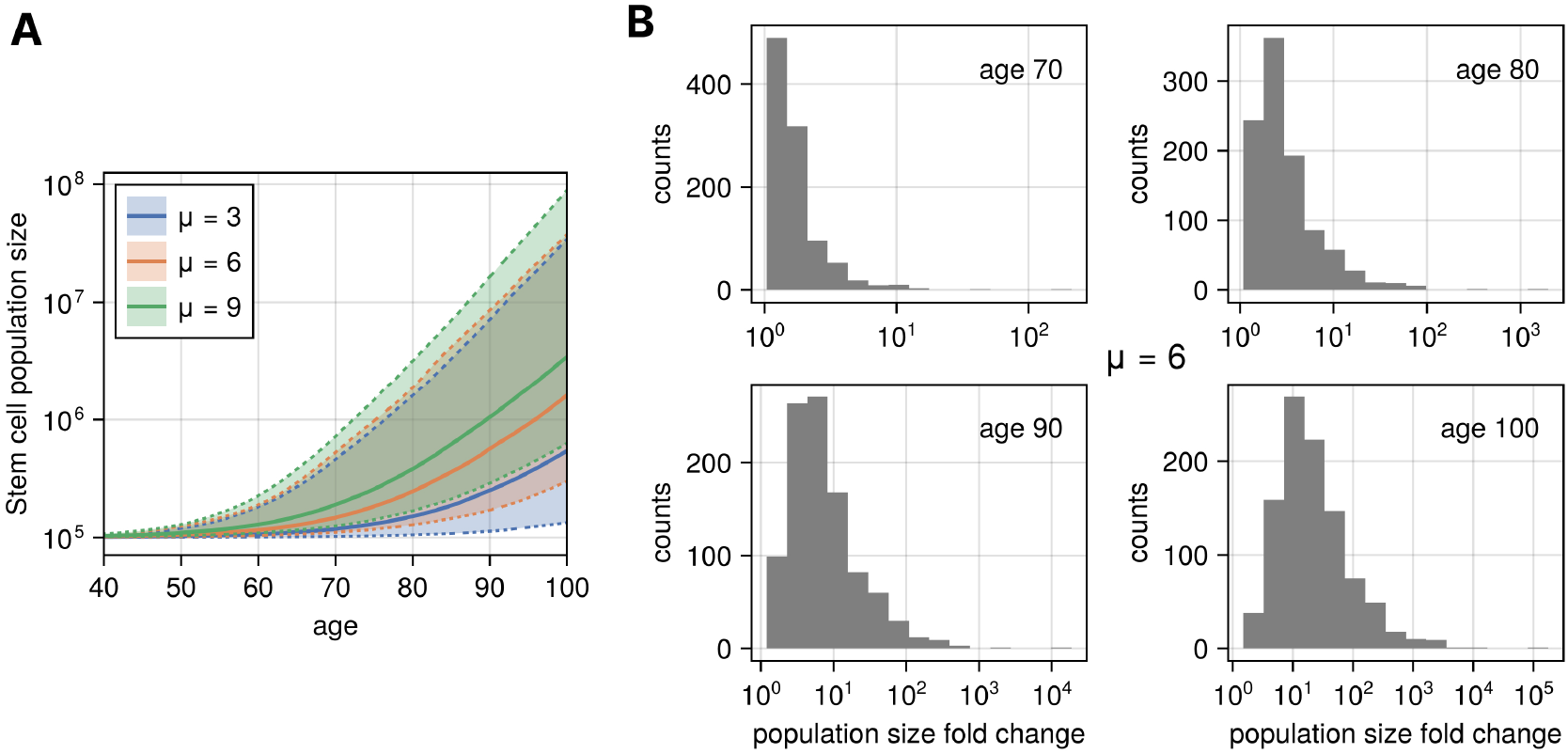
Unconstrained growth of clones leads to exponentially increasing stem cell populations. Under parameter values *η* = 0.08, *σ* = 0.025, *τ* = 1, and *N* = 10^5^, the HSC population increases many-fold from ages 70 to 100. **A** The median (solid lines) and 95% interval (shaded area) of the HSC population size across 1000 simulations for different values of the accumulation rate of fit clones *µ*. **B** The distribution of HSC population size fold changes for ages 70, 80, 90, and 100, obtained from 1000 simulations with *µ* = 6.

**Figure SI 15.**
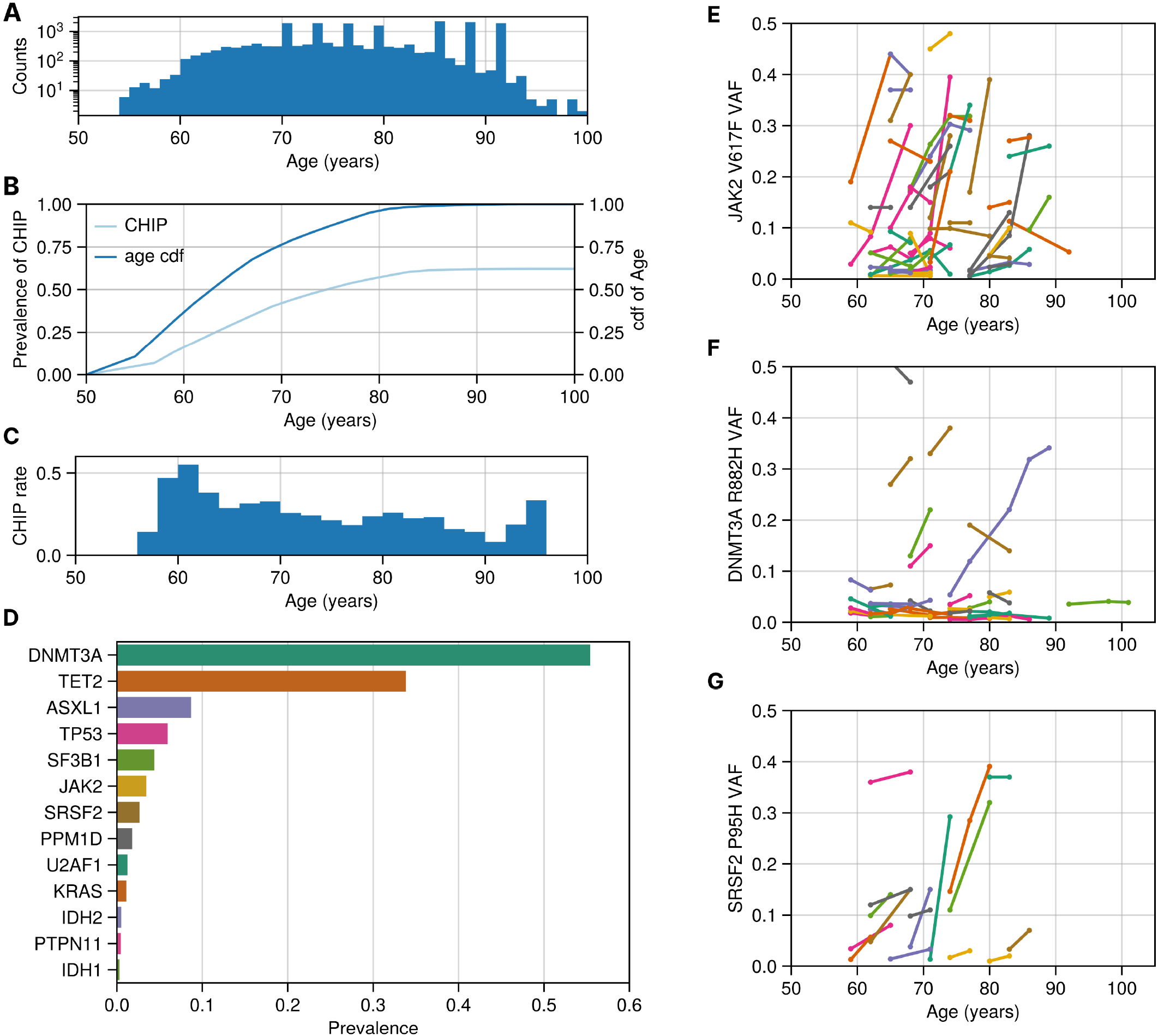
Aggregating three target sequencing datasets to study haematopoiesis in 1’923 individuals. **A**. The distribution of mutations per age. **B**. The prevalence of clonal haematopoiesis of indeterminate potential (CHIP) shown in light blue and computed as the cumulative sum of the prevalence of donors with at least one variant with VAF ≥ 2%. The cumulative distribution function (cdf) of the age of the donors (dark blue). **C**. The rate of CHIP computed by normalising the number of donors with CHIP within two-year age bins by the number of donors in each bin. **D**. Frequently mutated genes in this cohort. **E-G**. The dynamics of the VAF of three variants over time per different donors (each line represent an individual). *JAK2*-V617F was the most prevalent variant and was found in 67 donors, followed by *DNMT3A*-R882H which was found in 49 donors. *SRSF2*-P95H was found in 26 donors.

